# Segregating DNA lesions point to high selective advantage of tumor initiating cells

**DOI:** 10.1101/2025.10.02.680094

**Authors:** Vladimir Seplyarskiy, Maha Shady, Maria A Andrianova, Michael Spencer Chapman, Eliezer M Van Allen, Shamil R Sunyaev

## Abstract

The complications with identifying cells at the origin of cancer and tracking their early divisions impede studies of cancer initiation. Recently, it was shown that some DNA lesions generated by a pulse of damage-inducing mutagen persist over multiple rounds of replication. Segregation of DNA lesions in the early genealogy of an expanding clone leaves a statistically interpretable footprint of cancer initiating events. Specifically, it allows for estimating the number of cell divisions between the initiating DNA lesion and the most recent common ancestor of the tumor. Here, we analyze footprints of segregating lesions from a previously published experimental mouse system, as well as post-chemotherapy human metastatic tumors and the blood of chemotherapy treated patients. In all contexts, clones tend to start early, usually within the span of 4 cell generations from mutagen exposure. Using a branching process model, we show that fitness advantage of early cancer drivers exceeds 30%, with each early division leading to at least 1.3 self-renewing cells. We highlight an example of a blood-derived single cell phylogeny with major subclones separated by just two cell divisions. Broadly, our approach allows inference of tumor initiation and growth parameters based on events preceding the most recent common ancestor of the initiating clone as opposed to characteristics of fully grown tumors.

## Introduction

Cancer arises as a clonal expansion of a single cell. Most cancers start following the accumulation of driver mutations that result in a proliferative advantage of the clone^1^. Observations on the evolutionary dynamics of cancers (or clones in healthy tissues) include the distribution of clone sizes inferred from variant allele fractions^2–6^, driver mutation recurrence^7^ and, more recently, single cell phylogenies^8^. Although highly informative about neoplastic progression, these observations are influenced by long-term clone trajectories and lack resolution to provide insights into tumor initiation.

Recent studies have demonstrated that DNA lesions that persist unrepaired for multiple cell divisions could provide information about tumor initiation^9–12^. Aitken et al. used mutagen-induced mouse liver tumors to show that an experimentally initiated pulse of DEN mutagenesis creates DNA lesions that remain stable over multiple rounds of replication^9^. Successive replications over these lesions can generate different mutations at the same site resulting in multiallelic variants (MAVs) in the growing clone^9–12^. Due to random segregation of chromosomes during cell division, multiallelic variants would only exist in as many chromosomes as are inherited from the mutagenized strand by the most recent common ancestor (MRCA) of all tumor cells (Figure 1A,B). In absence of mitotic recombination, each division following mutagenesis but preceding the emergence of the tumor’s MRCA reduces the number of lesion-bearing chromosomes, and thus dilutes the number of chromosomes with detectable MAVs, on average, by twofold. Therefore, the system depicted in Figure 1 allows for estimating the number of cell divisions between the introduction of DNA lesions and the clonal expansion of the tumor based on the fraction of chromosomes carrying MAVs in the final tumor. We call this number “Lesion to Ancestor Divisions” (LAD) and develop a new statistical method for its estimation. We show that the distribution of LAD values among tumors is highly informative about early steps of cancer initiation and allows estimating important parameters such as the selective advantage conferred by the initiating driver mutations (Figure 1A).

**Figure 1.**
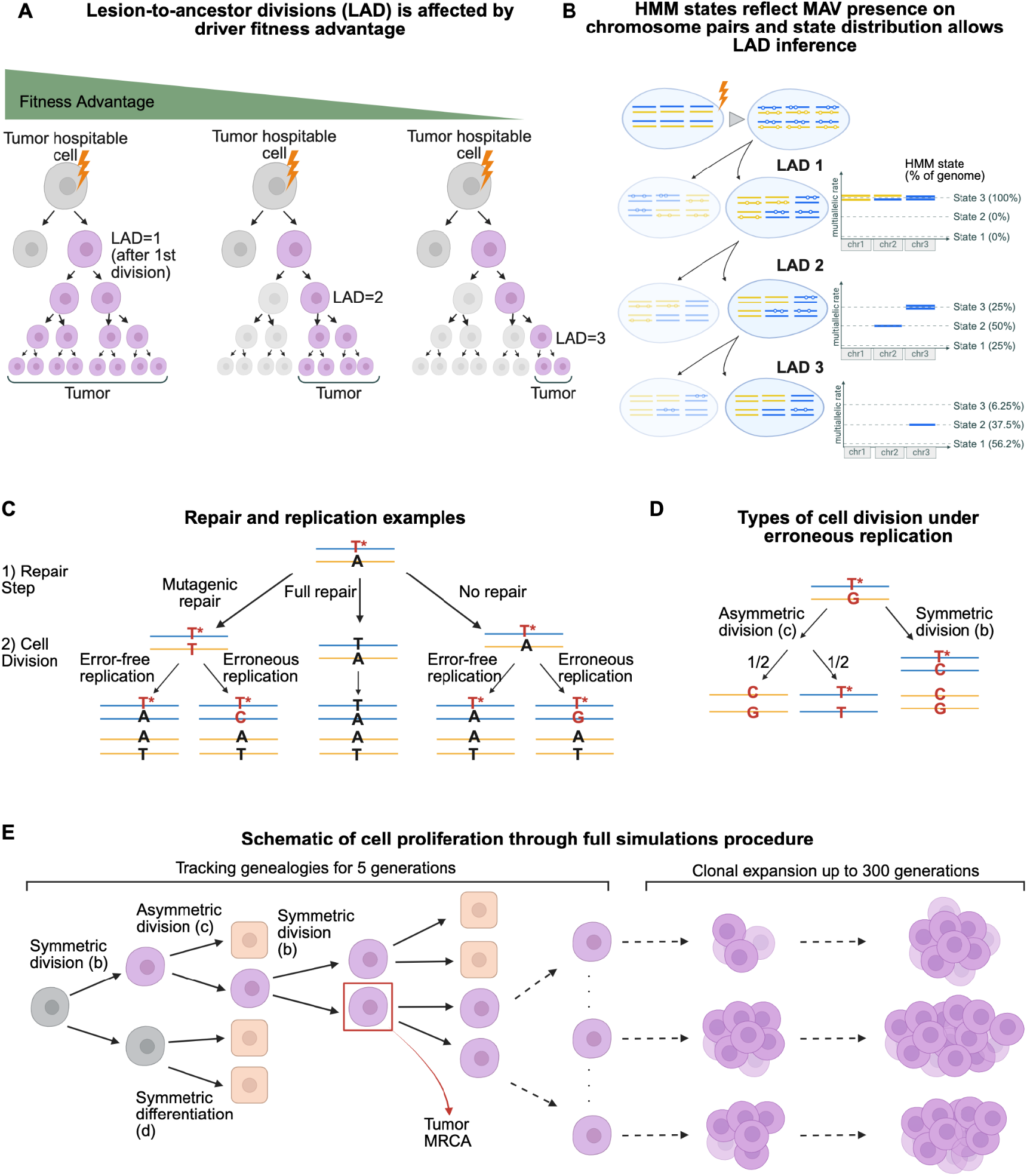
**A:** Schematic demonstrating how the fitness advantage of the acquired drivers affect the MRCA timing, with a higher fitness corresponding to an earlier MRCA. Purple cells carry driver mutation(s) or lesion-mutation heteroduplex(es). The cell that becomes the tumor MRCA must have a driver mutation on at least one strand at one locus, and it divides into two surviving lineages that contribute to the final tumor. **B:** Schematic of how chromosome segregation affects the number of lesion-bearing strands (marked with empty circles) at each division, and how that relates to the Hidden Markov Model (HMM) states used to estimate the LAD for each tumor (Methods). We show three pairs of homologous chromosomes to demonstrate that the first mutagenized cell carries lesions on all strands of all chromosomes, and that going through cell division with random segregation would yield daughter cells where both pairs of all chromosomes carry lesions (HMM state 3). Through another round of cell divisions, each daughter cell in each chromosome pair might inherit a lesion containing strand from both homologous chromosomes (HMM state 3), from only one (HMM state 2), or from neither (HMM state 1). **C:** Schematic of possible trajectories of a lesion through a full cycle of repair and replication. The nucleotides in red indicate a lesion or mutation, with the star specifying a lesion, and nucleotides in black indicate wild type. At each step, a lesion can undergo full repair (middle) with rate *r* − *u* leading to the wildtype nucleotides on both strands, and two wildtype daughter cells post division. Alternatively, a lesion can undergo mutagenic repair (left) with rate u, leading to a lesion-mutation heteroduplex, or it may not undergo repair (right) with rate 1 − *r*. For the cases where a lesion is maintained, replication over the lesion can be error-free with rate *ϵ* leading to the wildtype nucleotide inserted opposite the lesion, or erroneous with rate 1 − *ϵ* leading to a heteroduplex. **D:** Asymmetric division with erroneous replication (left) will lead to a stem cell with mutations on both strands half of the time, and a stem cell with a heteroduplex half of the time. Symmetric division (right) would lead to both stem daughter cells, one with a mutation and one with the heteroduplex. **E:** Schematic of the simulation procedure (Methods). Cell genealogies are tracked for at least 5 generations where cells undergo rounds of repair and division as described, square cells are differentiated and not tracked. After drivers are fixed, remaining stem cells are grouped into subclones based on genealogy, and they undergo clonal expansion and competition as described in the Methods. This figure was created in BioRender. Miler-jones, L. (2025) https://BioRender.com/rd257e4

In this system, a tumor forms because one or more DNA lesions are converted into one or more driver mutation(s). Cells possessing the driver(s) proliferate with one of them becoming the MRCA of the growing clone. The MRCA is the most recent individual cell that is the ancestor of all lineages in the tumor. This means that the MRCA is the first cell that divides symmetrically with both daughters giving rise to long-term lineages that contribute to the grown tumor. If cells frequently divide symmetrically and the resulting daughter cells establish stable lineages, clonal expansion would start very early. The initial expected growth rate of early clones is their fitness which is determined by the excess of symmetric division rate over the rate of terminal differentiation (or cell death). Thus, an early emergence of MRCA (small LAD value) points to an immediate strong increase in fitness and a high initial growth rate, while a delayed MRCA occurrence (large LAD value) points to lower selection intensity.

To connect LAD and fitness advantage we develop a branching process model of cancer evolution^13^ using previously published data from a mutagenized mouse system^9^. While studying the mouse system allows describing early stages of tumor initiation at high resolution, our primary interest is to investigate whether the same dynamics are observable in humans in natural or clinically relevant conditions. We subsequently discover multiallelic variants in a subset of human metastatic tumors that were treated with chemotherapy^14^. Furthermore, we analyze single cell phylogenies of human blood cells from chemotherapy-treated patients, which allows us to place lesion segregation events on individual branches^12,15^.

Estimation of key parameters of microevolution is the purview of population genetics. Over decades, the field has been developing sophisticated inference methods based on the amount of variation, frequency spectra, and shapes of genealogies. LAD distribution provides a completely new statistic that can be estimated from data in some cancers and other somatic clonal outgrowths. Unlike the analysis of genealogies or allele frequencies, this statistic is informative about the events between the introduction of a driver lesion and the emergence of the tumor MRCA.

## Results

### Estimating initial tumor growth rate from the LAD distribution

Cell divisions can lead to three outcomes: 1) a symmetric division resulting in two stem daughter cells, 2) an asymmetric division resulting in one stem cell and one differentiated cell unable to divide further, and 3) a symmetric division leading to two differentiated cells. Given that we only focus on the growing clone, we do not distinguish cell death and differentiation, as we assume that only stem cells contribute to the developing tumor. Prior to tumor initiation, the number of cells is kept constant. Thus, the rate of symmetric divisions resulting in two stem cells (birth rate) should be equal to that of symmetric divisions resulting in two differentiated cells (death rate). The rate of asymmetric divisions does not change the size of the cell population. Following tumor initiation, birth rate per generation (denoted ***b***) exceeds death rate per generation (denoted ***d***), and the expected tumor growth per generation is determined by the difference between birth and death rates (this rate equals ***1+b-d***). Thus, after acquiring driver mutation(s), growth depends on the selection coefficient of driver(s) ***s=b-d***.

We hypothesize that the time it takes to establish the tumor MRCA after lesion acquisition at driver loci is informative about the initial average rate of clonal growth (selection coefficient). As discussed, the MRCA is the first cell that divides symmetrically and produces two daughter lineages that both survive and proliferate long-term, leading to the final tumor (Figure 1A). We expect that a cell that acquires driver(s) which confer a high selective advantage is more likely to divide symmetrically, resulting in daughters that start surviving lineages and ultimately contribute to the developed tumor. As an extreme hypothetical example, a cell carrying driver(s) with ***b=1*** (implying ***s=1***) always produces two surviving lineages. The time (measured in cell divisions) it takes to establish a tumor MRCA after acquisition of driver lesion(s) that confer a low selective advantage is, on average, longer because more cell divisions are expected to either be asymmetric or to generate lineages destined for extinction.

Based on these principles, we develop a model of the tumor initiation and growth that incorporates cell divisions prior to and following the driver acquisition, the rate of DNA repair (characterized by parameter ***r***), and the rate of mutations arising from replication over lesions (parametrized by ***1-ε***). As noted in Anderson *et. al*^11^, a small fraction of mutations arise from lesions outside the S-phase, likely due to DNA synthesis opposite to the lesion during repair. We incorporate this process in the model (denoted by mutagenic repair with rate ***u***). We implement this model in computer simulations and, in the case of a single driver event, develop a straightforward analytical approximation to evaluate the distribution of LAD values and infer the initial clone growth rate (Methods; Figure 1C,D,E).

An interesting property of lesion segregation is that the MRCA does not necessarily possess the driver mutation on both DNA strands. It may contain a heteroduplex of the lesion opposite the driver mutation. Cell division of such an MRCA would generate one daughter cell carrying the driver mutation, and another carrying the lesion that will be converted to the driver at this or successive generations.

Remarkably, our calculations show that the LAD distribution depends on the compound parameter ***s=b-d***, which is the selection coefficient and the initial growth rate, and does not depend on other cell division parameters. Specifically, conditioning on ***s***, the LAD distribution does not depend on relative rates of symmetric and asymmetric divisions before and after the driver acquisition (Methods). This makes the LAD a key statistic for inferring ***s***. The LAD distribution does depend, however, on parameters related to DNA repair and translesion replication.

### Analysis of LAD distribution in chemically induced mouse liver tumors and inference of selection coefficients

Having established this conceptual framework for LAD inference, we subsequently analyze whole genome sequencing (WGS) data from 371 mouse liver tumors that were induced by a single pulse of the mutagen DEN and harvested 25 weeks later^9^. The tumors carry an average of 59,012 mutations per genome, nearly all of them caused by DEN^9^. DEN-induced mutations have highly specific and interpretable mutational profiles dominated by T>N and C>T substitutions. As convincingly shown by the original studies and noted above, DEN-induced pre-mutation lesions frequently persist for multiple cell divisions, undergoing translesion replication which leads to the incorporation of different mutations at the same position. Inspired by *Atkien et. al*.^9^, we develop a Hidden Markov Model (HMM) approach to estimate LAD using these multiallelic variants (MAVs), accounting for mitotic recombination^9^ (Methods). The approach relies on the following logic: the pulse of mutagen happened once in this system, so every cell division after the pulse decreases the fraction of chromosomes that carry DEN-induced lesions by a factor of two. As a result, the fraction of genomic segments that have multiallelic variants in the tumor should be proportional to *2*^*-n*^, where *n* is equal to LAD (Methods; Figure 1B).

The authors of the original study identified a known driver mutation in almost all (98%) DEN-induced liver tumors. In total, there are four recurrent driver mutations across 3 genes (two different mutations at the same site of *Hras*, one mutation in *Braf*, and one in *Egfr*). The driver mutations are mutually exclusive in 90% of tumors (excluding samples that likely represent a mixture of independent tumors; Methods). We excluded samples with more than one known driver from the analysis.

Since the LAD distribution depends on fitness advantage associated with driver mutations, we expect tumors with different drivers to have distinct distributions. We stratify tumor samples by known drivers, and estimate LAD distributions separately for each set of tumors. LAD distributions are indeed different between tumors driven by distinct drivers (Figure 2A,B,C; p<0.01 for *Egfr* 11:14185624_T/A vs *Hras* 7:145859242_T/A). Still, they share common features, most notably, more than 50% of tumors have MRCA emerging at the first or second cell divisions.

**Figure 2.**
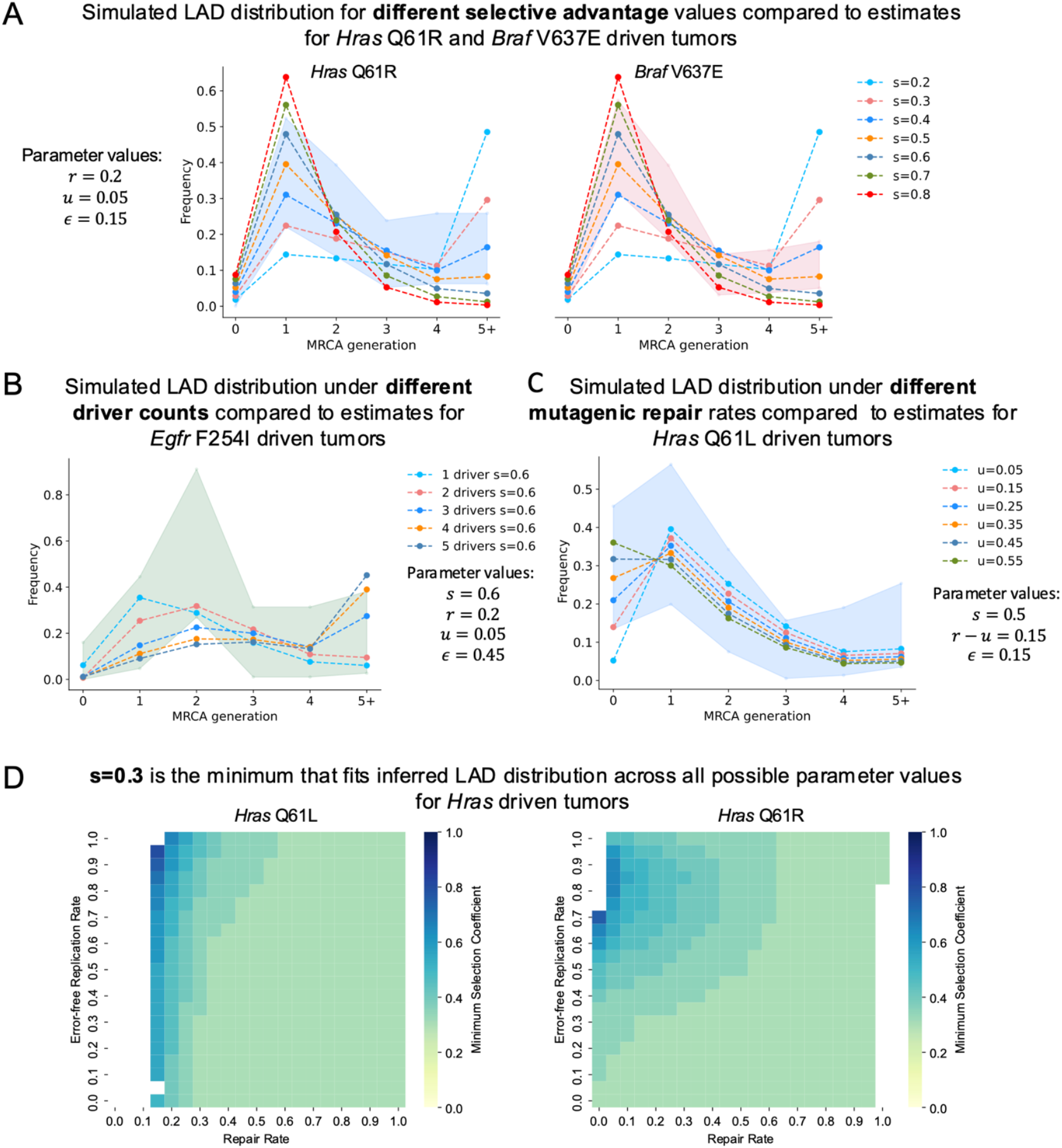
**A-C:** Simulated LAD distributions under different parameter combinations (shown by lines) against estimated LAD distributions (95% Poisson confidence intervals shown by shaded background) for each driver. **A** shows the simulated distributions for selection coefficient values 0.2-0.8 for a single driver under repair rate (*r*) of 0.2, mutagenic repair (*u*) of 0.05, and error-free replication rate (*ϵ*) of 0.15 against the estimates for *Hras* Q61R tumors (left) and *Braf* V637E tumors (right). **B** shows the simulated LAD distributions for 1-5 drivers, all with final selective advantage (*s*) of 0.6 under parameters *r* = 0.2, *u* = 0.05, *ϵ* = 0.45 against the estimates for *Egfr* F254I tumors. **C** shows the simulated LAD distributions with varying values of mutagenic repair (*u* = 0.05 − 0.55) and the remaining parameters *s* = 0.6, *r* − *u* = 0.15, *ϵ* = 0.15 against the estimates for *Hras* Q61L driven tumors. **D:** Minimum driver selective advantage that can fit the estimated LAD distribution for the *Hras* driven tumors (Q61L on the left and Q61R on the right) at each parameter combination. The y-axis of each heatmap represents error-free replication rate (*ϵ*), and the x-axis represents overall repair rate (*r*, mutagenic+full repair), the color at each box shows the minimum possible value of *s* over all possible values of *u* for this *ϵ, r* pair that can fit the estimates for the indicated driver assuming Poisson error.

We compute analytical approximations based on the branching process model we developed for all possible combinations of parameters (***s, ε, r*** and ***u***) in the single driver case, as well as run simulations for a subset of the possible parameter combinations in the single driver and multiple driver cases. We compare the model results with the observed LAD distributions assuming Poisson confidence intervals. First, all LAD distributions are consistent with the possibility of a single driver mutation resulting in the tumor growth. Interestingly, a broad range of DNA repair parameters that agree with empirical estimates from data can fit the LAD distribution for each driver (Methods). In contrast, for any parameter combination, the average tumor growth rate (selection coefficient ***s***) has to be at least 0.3 (Figure 2A,D; Supplementary Figures 1,2,3). We also find that fitting the LAD distribution inferred for *Hras* Q61L driven tumors, which are enriched for LAD=0 tumors, requires a higher value of mutagenic repair rate (***u***), suggesting that this process may contribute to an even earlier MRCA (Figure 2C; Supplementary Figures 1,2,3).

The model relies on the assumption that the cell carrying a driver mutation-lesion heteroduplex possesses fitness advantage. We demonstrate robustness of selection inference from LAD distribution with respect to this assumption using simulations (Methods).

Consistency of LAD distributions with the action of a single driver (as is almost always detected in the data) does not exclude the possibility of additional genetic or epigenetic driver mutations. We are able to fit models with two (albeit not more) drivers to LAD distributions of tumors with all 4 known drivers (Figure 2B, Supplementary Figures 3,4). In all simulations involving two drivers, the resulting fitness associated with the two drivers jointly is also very high (***s*** ≥ 0.3; Supplementary Figure 3), suggesting robustness of the parameter inference to the number of driver events.

**Figure 3.**
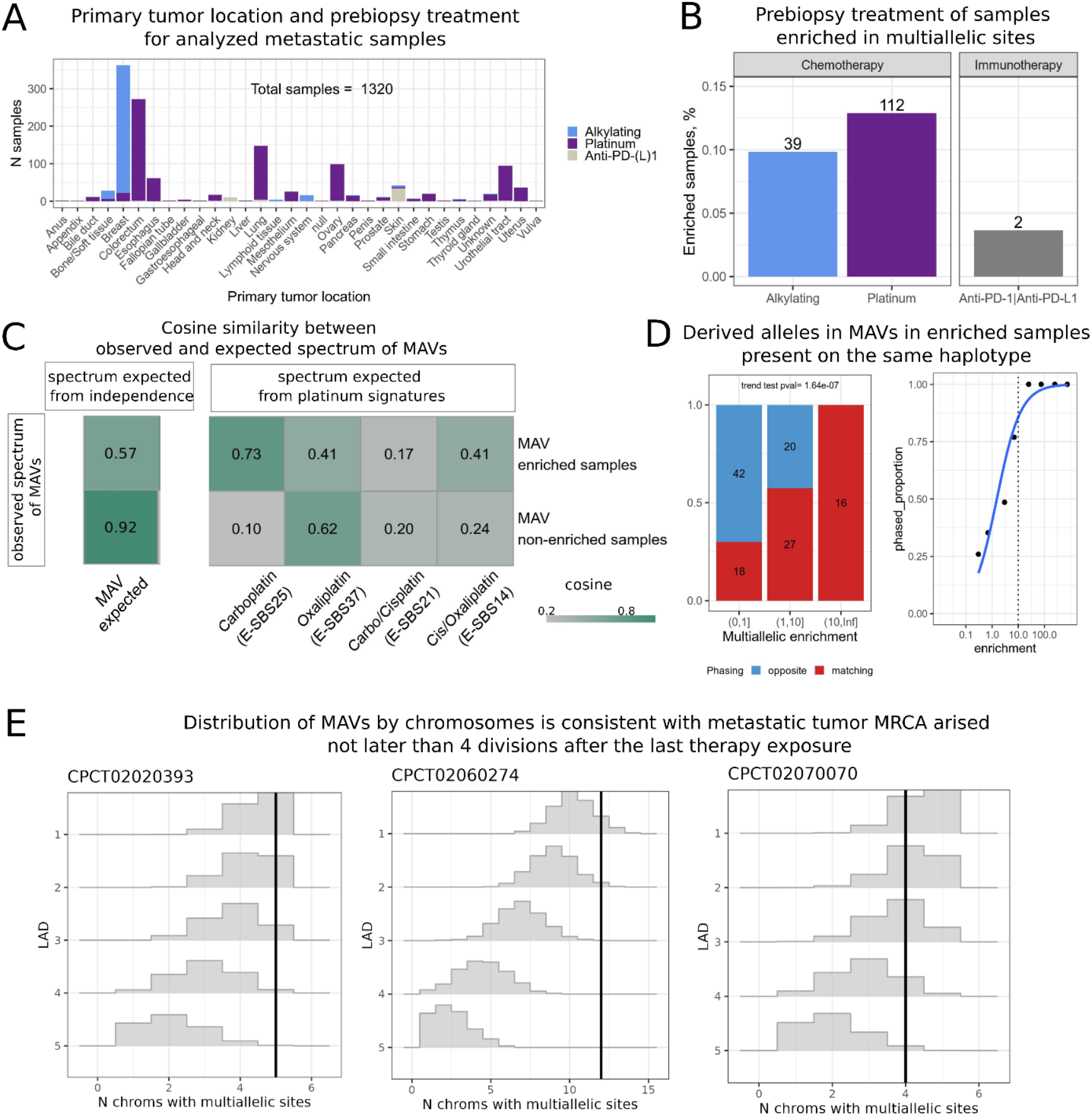
**A:** Distribution of analyzed metastatic tumors by location of primary tumor and prebiopsy treatment. **B**: Proportion of samples enriched in multiallelic sites in each prebiopsy treatment group. The number on the top of each bar represents the absolute number of the enriched samples. **C:** cosine similarities of MAVs spectra from enriched and non-enriched samples with the spectra expected for multiple independent mutations in the same site (left) and known spectra associated with different types of platinum treatment (right). **D:** Proportion of matching and opposite cases among phased multiallelic variants depending on the multiallelic enrichment. On the left, samples are split in 3 categories based on enrichment, numbers on the bars represent absolute values of the MAVs, p-value corresponds to the Armitage trend test. On the right, samples are split into more enrichment categories to show monotonic behavior of the function. **E:** Simulated distribution of the expected number of chromosomes containing multiallelic sites for three random tumors with 5+ MAVs as a function of LAD and the total number of observed MAVs. Black vertical line represents the real observed number of chromosomes with MAVs in the sample.

### Lesion segregation in human post-chemotherapy metastatic samples

Beyond experimental systems, pulses of mutagen are relevant for understanding oncogenesis in humans. It is established that chemotherapy treatment of primary tumors is associated with a high risk of secondary tumors and leads to the accumulation of mutations in metastatic clones^14– 18^. We hypothesize that multiallelic variants from post-chemotherapy metastatic tumors are informative about the fitness advantage of metastatic drivers. We thus reanalyze whole genome sequencing data from metastatic tumors aggregated by the Hartwig consortium^14^ and select 1320 samples across different types of primary cancer that were either exposed to known lesion-inducing mutagens: platinum (n=869) and alkylating agents (n=396), or to immunotherapy (n=55) treatment which we use as a control (Figure 3A).

Similarly to the experimental system discussed above, if mutagen exposure leads to segregating lesions, rounds of consecutive translesion replications would generate MAVs. Alternatively, MAVs may occur as independent mutations at the same site. MAVs originating from lesion segregation and MAVs due to independent mutations manifest differently in the sequencing data in terms of frequency, spectrum, and phasing. First, lesion segregation leads to an increase in the number of MAVs relative to the expectation based on the probability of independently co-occurring mutations. We find that almost all samples with over 10-fold excess MAVs come from patients on cytotoxic therapy but not on immunotherapy, consistent with DNA lesion persistence as a potential source of MAVs (Figure 3B). Note that we cannot rule out tumor type as a confounder because treatment strategy is strongly associated with tumor type.

Second, a MAV resulting from a chemotherapy-associated lesion should conform with the expected chemotherapy signature for the pair of mutations forming the MAV^12^. In contrast, MAVs formed by independently occurring mutations would be determined by the mutational profile of the tumor that is formed by multiple signatures. In platinum-treated patients, the spectrum of multi-allelic variants is more consistent with the carboplatin-associated signature E-SBS25 (we used signatures from Pich et al.^16^) than with the spectrum expected for independently co-occurring mutations (Methods), suggesting carboplatin as a source of persistent lesions leading to MAVs (Figure 3C).

Finally, MAVs originating from lesion segregation would appear on the same strand, and therefore phase to the same allele, whereas independent mutation events have at most an equal chance of being on the same or different alleles. For statistical reasons, two independent mutations occurring on the same allele would have a lower than equal chance to be detected (Methods). We phase derived alleles contributing to MAVs using neighboring germline SNPs. As expected under the lesion segregation model, all 16 phased mutations in tumors with signature E-SBS25 and excess MAVs above what is expected by independent mutations appear on the same allele (Chisq p-value <0.0001 for comparison with other tumors, Figure 3D). These results are consistent with the model where most MAVs with signature E-SBS25 originate from persistent lesions and are incompatible with independently occurring mutations or artifacts.

Unlike the experimental mouse system, human metastatic tumors carry many fewer MAVs (1-15 MAVs per tumor, with median 1 compared to 3756 in mice). This prevents a reliable estimation of the LAD distribution and corresponding model fitting. Still, the same logic is applicable: the number of chromosomes that carry lesions leading to MAVs decreases by a factor of two every division. Therefore, the distribution of MAVs across chromosomes can be used to estimate an upper bound on LAD. We simulate the expected number of chromosomes with MAVs for different LADs, accounting for the number of MAVs in a sample and for chromosome-specific mutability. The simulations suggest that for all 5 samples with 5 or more MAVs, the MRCA was established within less than 4 divisions following chemotherapy treatment (Figure 3E). As with observations in the mouse model, the LAD distribution dominated by an early appearance of MRCA points to a high fitness advantage of the growing metastatic clones. Given that we censor data points with fewer than 5 MAVs, which likely represent slower growing tumors, our inference of fitness advantage may be inflated when extrapolated to all metastatic samples.

### Evidence of rapid clonal expansions from single cell derived blood colonies

A series of recent studies have generated whole genome sequencing data from multiple single cell-derived colonies of human blood^8,12,15,19,20^. These data enabled the construction of underlying phylogenies with each somatic mutation placed on a specific phylogenetic branch. It has also been shown that lesion segregation could be deduced from the pattern of mutations on the phylogeny^12^. First, MAVs can manifest as a pair of different mutations at the same site on distinct nodes. Second, lesions persistent over a few generations that went through error-prone and error-free replications can generate the same mutations on different nodes; these are called PVVs (phylogeny violating variants) in the original work^12^ and they are impossible to identify without phylogenetically resolved data (Figure 4A). Both MAVs and PVVs suggest the presence of corresponding lesions on the ancestral node.

**Figure 4.**
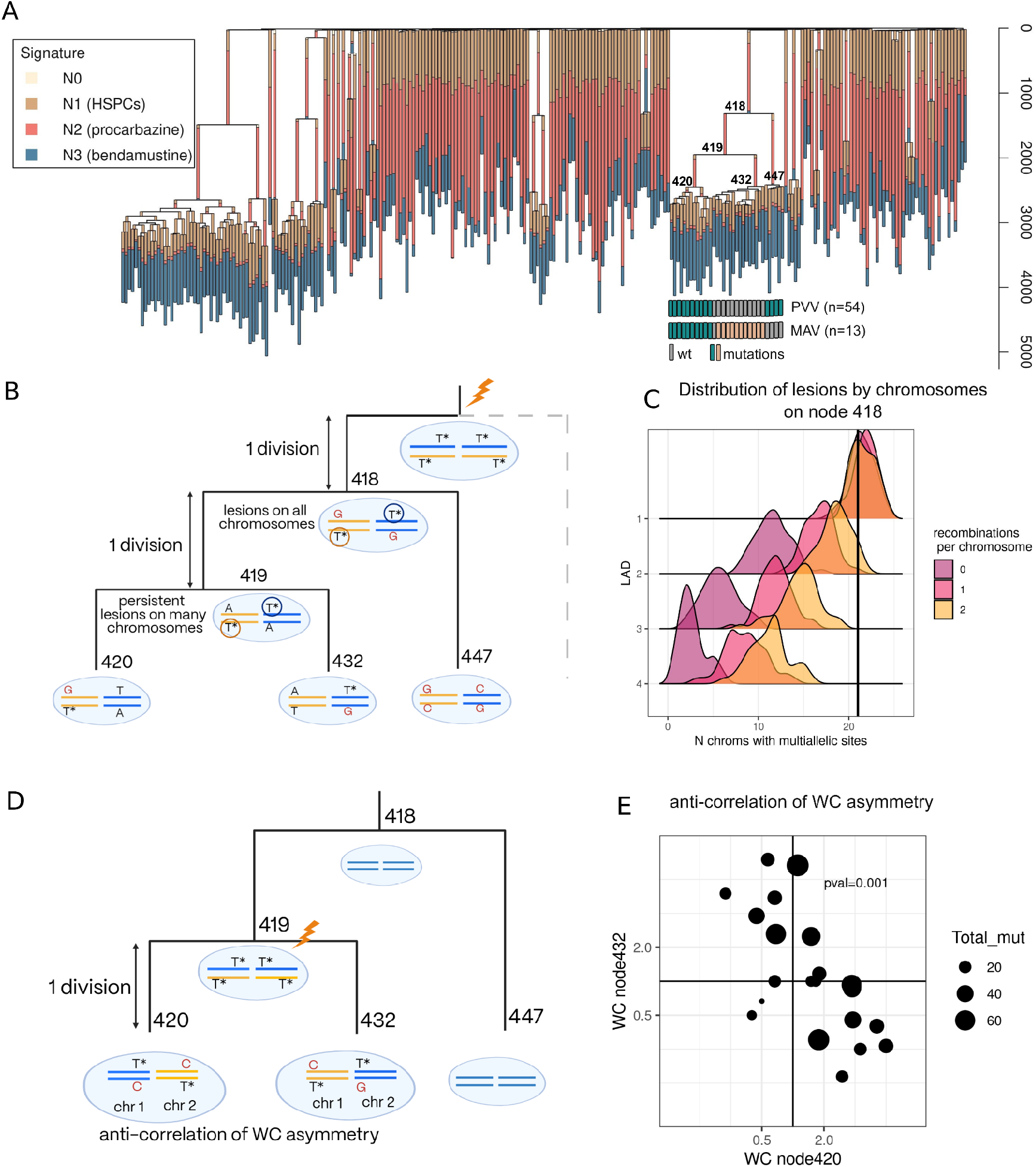
**A:** Phylogenetic tree of the single-cell expanded colonies from chemotherapy treated patient PX001_2_01. Branches are colored by the proportion of mutations on the branch attributed to the corresponding mutational signatures. Nodes of the clade with high amounts of MAVs and PVVs observed are enumerated for the reference in the text. **B:** Schematic explaining why PVVs spreading in many chromosomes imply single division from the pulse of chemotherapy to node 418 and a single division between nodes 418 and 419. **C:** Expected number of chromosomes containing PVVs on node 418 depending on different LAD and mitosis recombination rate. The vertical line corresponds to the observed number of chromosomes. **D:** Schematic representing mutational pattern expected in the case of single division between node 419 and node 420 (and 432): anti-correlation of Watson-Crick asymmetry (chromosome-level strand bias) between mutations attributed to node 420 and node 432. **E:** Anti-correlation of Watson-Crick asymmetry on the level of chromosomes between mutations attributed to node 420 and node 432 observed in the data (Spearman p-value=0.001).

We thus re-analyze data from two blood cancer patients treated with chemotherapy^12,15^. The treatment exposed their blood stem cells to pulses of mutagen, increasing the number of lesion-induced mutations. A substantial number of mutations that originate through lesion segregation was only detected in one of the patients, a female (PX001_2_01) treated with procarbazine for Hodgkin’s lymphoma, and focus our analysis on this patient. The patient’s available clinical history reports 6 cycles of ChIVPP (protocol containing procarbazine) chemotherapy at the age of 10 years for primary cancer treatment and 6 cycles of BGeV at the age of 47 years after relapse. Sequencing of 259 blood cell colonies from this patient reveals multiple clonal expansions (Figure 4). Overlaying mutation signatures over the phylogenetic tree topology suggests that the clonal expansions happened after procarbazine pulses at the age of 10 years.

For nodes with multiple MAVs and/or PVVs we can estimate how many divisions separate the pulse of chemotherapy and expansion of the clone. We detect 10 nodes in 9 independent subclades of the phylogeny with 5 or more MAVs+PVVs (Supplementary Table 2). Eight of them also have 5 or more chromosomes with at least one lesion (as marked by MAV or PVV) implying that lengths of branches stemming from these nodes are shorter than 4 divisions. In other words, fewer than 4 divisions separate two subsequent nodes on the tree.

Noticeably, node 418 (Figure 4A, B) has 13 MAVs and 54 PVVs distributed across 22 chromosomes. We account for the effect of mitotic recombination and conclude that exactly one division occurred between a pulse of chemotherapy and the branching point represented by node 418 (Figure 4C, Methods). Although the distribution of MAVs across chromosomes is not informative about the length of the subsequent branch (number of generations between nodes 418 and 419), the distribution of PVVs provides evidence that this branch also represents a single division. We can arrive at this estimate because the observed prevalence of PVVs across chromosomes requires persistence of lesions on daughter node 419, making it unlikely that the branch spans more than one division (Figure 4B).

An independent statistical footprint can be used to detect cases where a single division separates two sister nodes immediately after a mutagenic pulse. Two complementary DNA strands that acquire lesions in a single pulse will be inherited by the two daughter cells of the mutagenized cell. A lesion-containing strand in one sister will be mirrored by its complementary copy in the other sister cell (Figure 4D,E). As a result, if lesions lead to T>A mutations, a chromosome will be enriched in T>A mutations (polarized by reference strand) in progeny of one of the sister cells and in complementary A>T mutations (polarized by reference strand) in progeny of the other sister cell (Figure 4D,E). This will result in a broad pattern of anticorrelation in Watson-Crick pairing. This anticorrelation can be used to identify pairs of cells (nodes on the phylogenetic tree) separated from their damage-exposed common ancestor cell (the preceding node) by a single division.

Remarkably, we notice a strong anticorrelation in chromosome-level mutational Watson-Crick asymmetry between branches 419-420 and 419-432 (Figure 4E, spearman correlation p-value=0.01). This suggests that there is a subsequent pulse of chemotherapy after the one we reported for node 418. Moreover, we infer that the two internal phylogeny branches starting at node 420 and 432 are separated by a single cell division. Collectively, these observations paint a picture of an extraordinarily fast clonal growth triggered by two consecutive procarbazine pulses. These four nodes of the tree are separated by just two cell divisions (Figure 4B).

Intriguingly, we can estimate physical time spanning the generation preceding node 418 and the generation along the branch between nodes 418 and 419. As noted above, these two cell generations connect the penultimate and final pulse of procarbazine, which is usually administered through ChlVPP cycles separated by 2 weeks. This further demonstrates the potential of our lesion segregation analysis to measure key parameters related to early events in tumor development that otherwise remain obscure.

### PVVs are present in phylogenies of normal blood and mark nodes prone to expansions

Interestingly, PVVs and MAVs have been detected in single-cell derived colonies from normal blood that was not affected by chemotherapy or other treatment^12^. As we mention above, the probability for a pair of nodes to share the same lesion should decrease by at least a factor of two for every division that separates these nodes because of random segregation of lesion bearing chromosomes between sister cells. PVVs require the same lesion to be present in two consecutive nodes in the phylogeny, implying that these two nodes are unlikely to be separated by many cell divisions. Therefore, PVVs, on average, mark branches with small numbers of divisions. As we note above, a rapid start of the clonal growth is a marker of fitness advantage. A more intuitive and familiar footprint of positive selection is given by the overall number of progeny of a cell. We investigated whether these two footprints of rapid positive selection are in agreement with our intuition.

We calculate the number of branching events downstream of lesion nodes (Figure 5A) and compare it to the number of branching events downstream of a control set of nodes that have similar properties on a tree and are selected from the same individual (Methods). PVV-associated nodes have more downstream branching points, such that the number of lesion nodes with at least 10 downstream branching points exceeds the number of such control nodes by 2-fold (Figure 5B). We also compute the distance (measured in number of mutations) between nodes marking lesion segregation, which is the distance between a lesion node and a lesion repair node that define a PVV (Figure 5C). We find that the distance is more than 2 times lower for lesion segregation nodes compared to random pairs of control nodes with the same topology (Figure 5 D). These two observations suggest that the presence of PVVs is indeed a marker of high fitness advantage.

**Figure 5.**
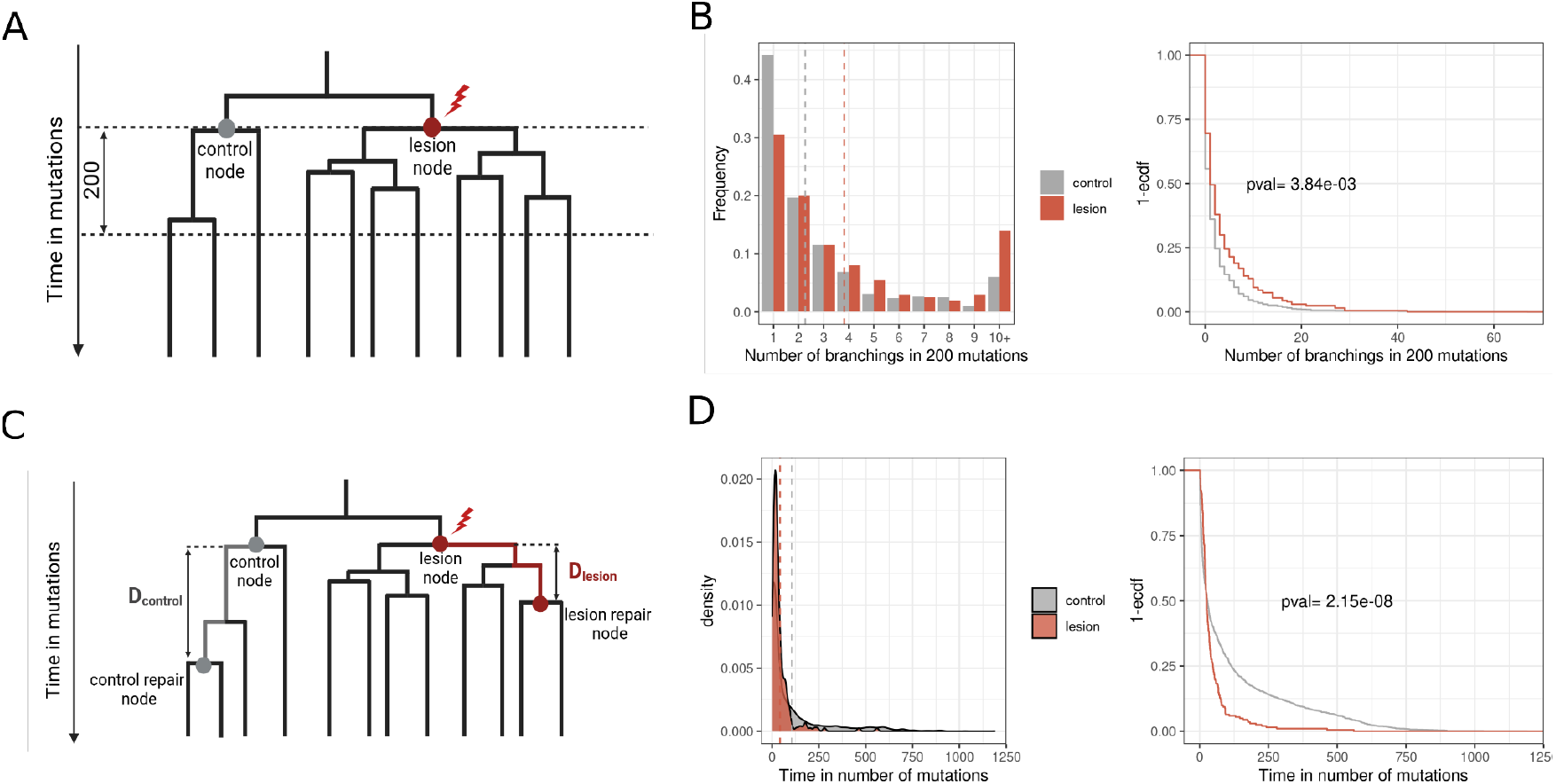
**A:** Schematic representing comparison of the number of branching points in a fixed time after node with lesion and control node (without lesion). **B:** Empirical difference in normal blood for statistic described in **A. C:** Schematic representing comparison of time (in mutations) between lesion node and node of lesion repair with the comparable control nodes (without lesion). **D:** Empirical difference in normal blood for the statistic described in **C**.

## Discussion

The phenomenon of lesion segregation offers insight into very early stages of somatic clonal expansion. Pioneering studies on mouse models showed that segregating lesions leave statistical footprints informative about the time span between the introduction of DNA damage and the establishment of the tumor’s MRCA. We show that the distribution of such times (LAD) primarily depends on the selection coefficient of driver mutation(s). Consequently, the minimal selection coefficient can be estimated from the LAD distribution under a broad range of assumptions about other relevant parameters. This selection coefficient is identical to the initial expected growth rate of the clone. The analysis suggests that DEN-induced liver tumors in mice have a very high initial growth rate increasing by at least 30% every generation.

Presence of segregating lesions are not limited to artificial systems. They were observed in primary cancers^9^, in post-chemotherapy human metastatic tumors^15^, and are even detected at low levels in untreated human blood^12^. Although the number of informative lesion-induced mutations is much smaller in natural human samples^12,14^, the basic conclusion of the fast initial growth due to newly acquired driver mutation(s) that confer a high fitness advantage remains relevant for human tumors. New data on single cell derived colonies allow the detection of individual rapidly growing clones starting with segregating lesions. We detect an example of a large clone and a nested subclone separated by just one cell division.

Existing work on selection inference in somatic clonal growth primarily analyzes VAF distribution statistics related to long-term growth rate^21–24^. The analysis of lesion segregation offers new observables informative about the initial cell divisions leading to the creation of the clone^9^ rather than its long-term evolutionary trajectory. This analysis framework is underexplored in the field of human somatic clone evolution as well as in population genetics more broadly. Although existing theoretical work incorporates some features of our system, including the effect of mutations on the subsequent phylogeny^25^ and the occurrence of (relevant) mutations during cell divisions^26^, it does not consider lesion segregation and LAD.

This approach could help address additional questions related to the early stages of clonal expansion. For example, the accumulation of chemotherapy-treated metastatic samples could help address the long standing question of how chemotherapy stimulates clonal expansions: is it through the creation of new mutations, or by selecting pre-existing somatic variants or cells in specific transcriptional states? Clonal growth fueled by a new somatic driver is highly unlikely to start with LAD=0 (the cell exposed to the mutagen becoming the tumor MRCA) because the resulting daughter lineages will possess mostly non-overlapping sets of damage-induced mutations. On the contrary, for cells with pre-existing genetic or epigenetic driver(s), damage-induced mutations are irrelevant in terms of their fitness advantage. The prevalence of secondary tumors estimated to have LAD of 0 based on this methodology could inform the role of chemotherapy-induced mutations in driving secondary tumors.

## Acknowledgements

The authors would like to thank Johannes Berg, Nuria Lopez Bigas, and Matt Pennel for their insightful comments on this work, as well as Mikhail Moldovan for constructive discussions on estimating selection in post-chemotherapy metastatic tumors. This work was funded by the National Institutes of Health (NIH) National Human Genome Research Institute (NHGRI) grant U01HG012009 (to SRS) and NIH National Institute of General Medical Sciences (NIGMS) grant R35GM127131 (to SRS). MS was partially funded by the Eric and Wendy Schmidt Center (EWSC) at the Broad Institute of MIT and Harvard.

## Author Contributions

V.S., M.S., M.A.A., and S.R.S designed the study and wrote the manuscript. V.S. and M.A.A. developed and applied statistical analyses to the mouse and human data. M.S. and S.R.S. developed the branching process model and implemented the corresponding analytical calculations, and M.S implemented and analyzed the corresponding computer simulations. M.S.C assisted with the analysis of genealogies for single cell derived blood colonies. E.M.V.A provided guidance and feedback. All authors read, edited, and approved the final manuscript.

## References

1. Greaves, M. & Maley, C. C. Clonal evolution in cancer. Nature 481, 306–313 (2012).

2. Abby, E. et al. Notch1 mutations drive clonal expansion in normal esophageal epithelium but impair tumor growth. Nat. Genet. 55, 232–245 (2023).

3. Williams, M. J. et al. Quantification of subclonal selection in cancer from bulk sequencing data. Nat. Genet. 50, 895–903 (2018).

4. Yokoyama, A. et al. Age-related remodelling of oesophageal epithelia by mutated cancer drivers. Nature 565, 312–317 (2019).

5. Martincorena, I. et al. Somatic mutant clones colonize the human esophagus with age. Science 362, 911–917 (2018).

6. Körber, V. et al. Detecting and quantifying clonal selection in somatic stem cells. Nat. Genet. 57, 1718–1729 (2025).

7. Weghorn, D. & Sunyaev, S. Bayesian inference of negative and positive selection in human cancers. Nat. Genet. 49, 1785–1788 (2017).

8. Spencer Chapman, M. et al. Clonal selection of hematopoietic stem cells after gene therapy for sickle cell disease. Nat. Med. 29, 3175–3183 (2023).

9. Aitken, S. J. et al. Pervasive lesion segregation shapes cancer genome evolution. Nature 583, 265–270 (2020).

10. Aitken, S. J. et al. Genetic background sets the trajectory of cancer evolution. Preprint at 10.1101/2025.01.13.632787 (2025).

11. Anderson, C. J. et al. Strand-resolved mutagenicity of DNA damage and repair. Nature 630, 744–751 (2024).

12. Spencer Chapman, M. et al. Prolonged persistence of mutagenic DNA lesions in somatic cells. Nature 638, 729–738 (2025).

13. Kendall, D. G. Birth-and-Death Processes, and the Theory of Carcinogenesis. Biometrika 47, 13 (1960).

14. Priestley, P. et al. Pan-cancer whole-genome analyses of metastatic solid tumours. Nature 575, 210–216 (2019).

15. Mitchell, E. et al. The long-term effects of chemotherapy on normal blood cells. Nat. Genet. 57, 1684–1694 (2025).

16. Pich, O. et al. The mutational footprints of cancer therapies. Nat. Genet. 51, 1732–1740 (2019).

17. Pich, O. et al. The evolution of hematopoietic cells under cancer therapy. Nat. Commun. 12, 4803 (2021).

18. Liu, D. et al. Mutational patterns in chemotherapy resistant muscle-invasive bladder cancer. Nat. Commun. 8, 2193 (2017).

19. Mitchell, E. et al. Clonal dynamics of haematopoiesis across the human lifespan. Nature 606, 343–350 (2022).

20. Lee-Six, H. et al. Population dynamics of normal human blood inferred from somatic mutations. Nature 561, 473–478 (2018).

21. Poon, G. Y. P., Watson, C. J., Fisher, D. S. & Blundell, J. R. Synonymous mutations reveal genome-wide levels of positive selection in healthy tissues. Nat. Genet. 53, 1597–1605 (2021).

22. Hu, Z. et al. Quantitative evidence for early metastatic seeding in colorectal cancer. Nat. Genet. 51, 1113–1122 (2019).

23. Sun, R. et al. Between-Region Genetic Divergence Reflects the Mode and Tempo of Tumor Evolution. Nat. Genet. 49, 1015–1024 (2017).

24. Naxerova, K. Evolutionary paths towards metastasis. Nat. Rev. Cancer 25, 545–560 (2025).

25. Coupling adaptive molecular evolution to phylodynamics using fitness-dependent birth-death models | eLife. https://elifesciences.org/articles/45562.

26. Goldberg, E. E. & Igić, B. Tempo and Mode in Plant Breeding System Evolution. Evolution 66, 3701–3709 (2012).

