## Supplementary material for "Segregating DNA lesions point to high selective advantage of tumor initiating cells": Methods and Supplement

#### Simulations of tumor initiation and growth

##### Simulating cell

A mouse hepatic stem cell is initialized with 19 pairs of autosomes each carrying 5 randomly selected potential driver loci. This cell can then be used as the starting point for different iterations of lesion introduction and subsequent clonal expansion as will be described next.

##### Simulating Lesions

To simulate DEN exposure, lesions are randomly introduced at a specified number of these driver loci. Each lesion is randomly assigned to a strand. For simulations starting with a single driver lesion, the selection coefficient ($s_1$) of the introduced driver lesion is specified. For simulations with multiple driver lesions ($n$), we specify the selection coefficient of the first driver position ($s_1$) and the total potential multiplicative selective advantage of all the driver positions ($S_{max}$). The assigned selection coefficient of each of the remaining drivers is then set as $\sqrt[n-1]{\frac{1+s_{max}}{1+s_1}} - 1$. Note that since we are specifically looking at cell divisions, the maximum possible selection coefficient value is 1. Each set of introduced driver lesions along with their assigned selection coefficients can then be used to simulate the cell growth and clonal expansion process multiple times with different expansion parameters as will be described next.

##### Simulating cell growth and clonal expansion

After the lesion introduction step which simulates the mutagen exposure in mice, each simulation for the growth and expansion of the mutagenized cell consists of two stages. Stage 1 simulates rounds of lesion repair, fitness computation of each cell being tracked, and cell division. The process starts with a single mutagenized cell, and then tracks the descendants of that cell. This stage lasts for at least 10 generations and until all driver lesions are resolved by either turning into a mutation or being repaired. The detailed genealogy of each resulting cell is tracked for at least 5 generations or until all driver lesions are resolved. Stage 2 starts by grouping the resulting cells from stage 1 into subclones based on the genealogies as will be described later, and then simulates clonal expansion and competition of those clones based on each clone's fitness for up to 300 generations or until the total number of cells reaches $10^8$.

**Stage 1: Initial Stage of Tumor Growth**

Repair

At each generation, the lesions are first subject to a repair procedure. For each lesion, the repair process can 1) lead to a mutation being introduced opposite the lesion with rate $u$ (mutagenic repair), or 2) lead to full removal of the lesion with rate $r - u$. Each lesion has a chance of $1 - r$ of not being subject to repair at all.

Fitness computation

After the repair step, the fitness of each cell is computed to determine the type of cell division the cell will undergo. We assume that a driver contributes to a cell's fitness if both strands have a mutation at this locus or if there is a lesion opposite a mutation at this locus (lesion-mutation heteroduplex). We conducted experiments to evaluate the robustness of our results in the absence of this assumption. Specifically, we ran simulations for 112 parameter combinations either assuming that a driver only contributes to a cell's fitness if there is a mutation on both strands, or if presence of a lesion on that strand at the driver loci is enough for a fitness gain. We then assessed how often a change in this assumption would affect the parameter fits for each of the *Hras* drivers, and found that at most 6.25% of parameter combinations that fit the data under our assumption would not fit if we changed it.

We run simulations with the multiplicative fitness model as well as with synergistic epistasis. If a simulation is being run under the multiplicative fitness model, then the final fitness of a given cell is computed as $w = \prod_i (1 + s_i)$ for each contributing driver $i$. If a simulation is being run under synergistic epistasis, then a cell would only start to acquire a fitness advantage with the second driver, in other words, a cell with one driver has no fitness advantage. In this case, if there are at least 2 driver mutations, the cell's fitness is computed the same way as in the multiplicative case.

Division and strand replication

Each cell can divide symmetrically into two new stem cells (birth process, $b$) or into two differentiated cells (death process, $d$), or asymmetrically into one stem cell and one differentiated cell ($c$), such that $b + c + d = 1$. Through analytical calculations, we determined that the MRCA distribution of the resulting tumor with respect to the generation when all driver lesions have been resolved is dependent only on s, and does not vary with the asymmetric division rate. The calculations are based on the following assumptions:

1) The selective advantage of a cell ($s$), corresponds to its growth rate such that $s = b - d$
2) The probability of a cell's lineage going to extinction is $\frac{d}{b}$[1], which can be derived with the following computation. Let $\rho$ be the probability of extinction and note that the probability of stem cell's lineage going to extinction is the sum of the probabilities that 1) the stem cell divides to two stem cells and both of them go

extinct, 2) the cell divides asymmetrically to one differentiated cell and one stem cell that goes extinct and 3) the cell divides into two differentiated cells:

$$\rho = b\rho^2 + c\rho + d$$
$$0 = b\rho^2 + (c-1)\rho + d$$
$$0 = b\rho^2 + (1-b-d-1)\rho + d = b\rho^2 - (b+d)\rho + d$$
$$0 = (b\rho - d)(\rho - 1)$$
$$\rho = \frac{d}{b}, \quad if\ b > d$$

And thus the probability of a cell's lineage surviving long term is $1 - \rho = \frac{b-d}{b} = \frac{s}{b}$

Keeping these relationships in mind, we set the first generation when all driver lesions are resolved as generation 0. To get a tumor with a cell in that generation as the MRCA (LAD=0), that cell needs to undergo a birth event, with both daughter cells starting lineages that survive through later generations, conditioned on the original cell forming a surviving lineage long term. This is a probability of

$$P(LAD = 0) = b(\frac{s}{b})^2 \frac{1}{(s/b)} = s$$

Moving forward, to have a tumor with the MRCA cell in generation 1, we need to either: 1) have an asymmetric division leading to generation 1, followed by a birth event for the resulting stem cell with both of its daughter cells surviving, or 2) have a birth event, with one daughter's lineage going to extinction and the other one undergoing a birth event with both daughter cells surviving, all conditional on the original cell forming a surviving lineage long term:

$$P(LAD = 1) = cb(\frac{s}{b})^2 \frac{1}{s/b} + 2 * b(\frac{d}{b})b(\frac{s}{b})^2 \frac{1}{s\backslash b} = cs + 2ds = (2d + c)s$$
$$P(LAD = 1) = (2d + 1 - b - d)s = (d - b + 1)s = (1 - s)s$$

Following this logic for later generations, we find that the probability of having a tumor with MRCA in generation k, follows the trend:

$$P(LAD = k) = (1 - s)^k s$$

This is a geometric distribution parameterized by $s$, and is independent of the asymmetric division rate. We verified that indeed in simulations the LAD distribution following driver resolution does not vary with the asymmetric division rate and follows a geometric distribution parameterized by s.

As such, we set the asymmetric division rate ($c$) as a pre-determined input parameter in our simulations, and compute the birth and death rates of a cell with selective advantage $s$ as:

$$b = \frac{s + 1 - c}{2}, \quad d = 1 - b - c$$

In the simulations presented here, we set $c = 0$. We show later in the section titled "Analytical Approximation of LAD Distribution" that even considering the full process that starts with a lesion rather than a fixed driver mutation yields an LAD distribution that does not depend on c for the single driver case.

Once the type of the cell division is determined, the cell division process itself involves random chromosome segregation with error-free translesion replication occurring with probability $\epsilon$, and replication over a mutation always leading to a mutation.

Grouping cells into subclones

Cells resulting from this process are then grouped together into subclones based on shared genealogy. The cell genealogies are tracked for the first 5 generations, and for any branch with unresolved driver lesions at the 5th generation, it continues to be tracked until driver lesion resolution.

##### Stage 2: Clonal expansion

After grouping cells into subclones, we simulate clonal expansion and competition for 300 generations or until we reach $10^8$ cells, whichever is earlier. At this stage, to simulate the next generation, we draw cells from the current generation using a multinomial distribution parameterized by each subclone's prevalence and relative fitness in the current generation. The overall size of the tumor at each generation follows exponential growth with growth rate equal to the average fitness of all subclones in the previous generation. Let $n_{i,t}$ and $w_{i,t}$ be the size and fitness of subclone $i$ in generation $t$, and $N_t$ be the overall tumor size at generation $t$,

$$\overline{w_t} = \sum_i \frac{n_{i,t}}{N_t}$$

$$N_{t+1} = N_0 \, \overline{\mathrm{w_t}}^{(t+1)}$$

$$\{n_{1,t+1}, n_{2,t+1}, \dots\} \sim Multinomial(N_{t+1}, [p_{1,t+1}, p_{2,t+1}, \dots]), where\ p_{i,t+1} = \frac{n_{i,t}}{N_t}\frac{w_{i,t}}{\overline{\mathrm{w_t}}}$$

The final set of subclones from this procedure is considered to comprise the final tumor. We consider a detectable tumor to have formed and proceed with the downstream analysis if there are at least 1 million cells.

##### Computing LAD

We run the described simulation procedure many times (10-500k) for each set of parameters, and compute the LAD distribution for each set of parameters to compare to the distributions observed in data.

To compute the LAD from one simulation, we follow the genealogies for all subclones in the final tumor to find the earliest generation with surviving lineages from both branches. If the MRCA generation is found to be 5 or greater, the tumor is assigned LAD = '5+' (this is because the HMM LAD inference from data, described below, cannot distinguish between LAD values that are 5 or greater). If the MRCA generation is found to be earlier, then we implement a recursive filtering procedure to simulate the detection limit of small clones that would be represented in the data. Specifically, we assume that a subclone that represents less than 10% of the final tumor would carry mutations with variant allele frequencies too low to be detected in the sequenced tumor sample and thus will not be accounted for in the HMM LAD inference. In this procedure, if the LAD is determined to be k < 5, then we compute the total proportion of the descendants of the cell at generation k, and if one of them is less than 10% of the overall cell population size, the whole branch is removed and the MRCA is re-computed. We continue this procedure recursively, until we either reach an MRCA generation for which both resulting branches are at least 10% of the full tumor size, or we reach an MRCA generation of '5+'.

#### Analytical Approximation of LAD distribution

Here we analytically compute the expected LAD distribution using the same model as the simulation procedure in the single driver case. The four types of cells that we can have in this scenario with respect to the driver locus are: 1) a cell with no lesions or mutations (wildtype nucleotide on both strands), we call this a $ww$ cell 2) a cell carrying the driver lesion on one strand and the wild type nucleotide on the opposite strand, we call this an $lw$ cell, 3) a cell carrying the driver lesion with a mutation having been introduced opposite the lesion, we this call an $lm$ cell, and 4) a cell with a mutation having formed at the driver locus on both strands, we call this an $mm$ cell.

Recall that the simulation procedure consists of sequences of repair followed by division. Also recall that in our formulation the $lm$ and $mm$ cells would carry the selective advantage of the driver, while the $lw$ and $ww$ cells wouldn't. We start by considering the possible trajectories of each cell type when undergoing those procedures.

Repair:

1) The $lw$ cell can be repaired to an $lm$ cell with probability $u$, to a $ww$ cell with probability $r - u$, and can remain as an $lw$ cell with probability with $1 - r$
2) The $lm$ cell can be repaired to an $mm$ cell with probability $r$, or stay as an $lm$ cell with probability $1 - r$
3) The $mm$ and $ww$ cells would not be subject to repair

Cell division:

1) If the $lw$ cell undergoes a birth event, it will lead to a $ww$ cell resulting from the lesion free strand, and either an $lw$ or $lm$ cell resulting from the lesion containing strand depending on whether replication over the lesion is erroneous or not. Erroneous replication over the lesion would occur with a probability of $1 - \epsilon$ and would lead to an $lm$ cell, while error-free replication over the lesion would occur with a probability of $\epsilon$ and lead to an $lw$ cell. If the cell undergoes asymmetric

division, half the time the resulting stem cell will come from the lesion containing strand and half the time it would come from the lesion free strand. Overall this would lead to the following event probabilities resulting from an $lw$ cell division:

$$p(lw \text{ } and \text{ } ww \mid lw) = b\epsilon$$
$$p(lm \text{ } and \text{ } ww \mid lw) = b(1-\epsilon)$$
$$p(ww \mid lw) = 0.5c$$
$$p(lw \mid lw) = 0.5c\epsilon$$
$$p(lm \mid lw) = 0.5c(1-\epsilon)$$

2) Following the same logic for an $lm$ cell, we would end up with the following events:

$$p(lw \text{ } and \text{ } mm \mid lm) = b\epsilon$$
$$p(lm \text{ } and \text{ } mm \mid lm) = b(1-\epsilon)$$
$$p(mm \mid lm) = 0.5c$$
$$p(lw \mid lm) = 0.5c\epsilon$$
$$p(lm \mid lm) = 0.5c(1-\epsilon)$$

3) Division of an $mm$ cell would lead to 2 $mm$ cells with a probability $b$ and one $mm$ cell with a probability $c$

$$p(mm \text{ } and \text{ } mm \mid mm) = b$$
$$p(mm \mid mm) = c$$

4) Division of a $ww$ cell would always lead to 1 or more $ww$ cells which cannot contribute to the final tumor so we ignore trajectories involving division of a $ww$ cell in our calculations, and ignore the $ww$ cell lineage in trajectories that lead to a $ww$ cell along with another cell

Putting the repair and cell division steps together, we get the following trajectories through 1 generation starting from each cell type. Notice here we use the initial birth and asymmetric division rates ($b_0$ and $c_0$) when starting with an $lw$ cell as this cell type would not have a fitness advantage according to our assumptions.

1) Starting from an $lw$ cell:

$$p(lw \mid lw) = u(0.5c)\epsilon + (1-r)(b_0 + 0.5c_0)\epsilon$$

Notice that since $b_0 + c_0 + d_0 = 1$, and we consider the initial cell population to be in equilibrium ($b_0 = d_0$), we can write $c_0 = 1 - 2b_0$, or $b_0 + 0.5c_0 = 0.5$, and thus:

$$p(lw \mid lw) = 0.5uc\epsilon + 0.5(1-r)\epsilon = 0.5\epsilon(uc + 1 - r)$$
$$p(lm \mid lw) = 0.5uc(1-\epsilon) + 0.5(1-r)(1-\epsilon) = 0.5(1-\epsilon)(uc + 1 - r)$$
$$p(mm \mid lw) = 0.5uc$$
$$p(lw \text{ } and \text{ } mm \mid lw) = ub\epsilon$$
$$p(lm \text{ } and \text{ } mm \mid lw) = ub(1-\epsilon)$$

2) Starting from an $lm$ cell:

$$p(lw \mid lm) = 0.5c(1-r)\epsilon$$

$p(lm \mid lm) \ = \ 0.5c(1-r)(1-\epsilon)$
$p(mm \mid lm) \ = \ rc + (1-r)(0.5c)$
$p(lw \ and \ mm \mid lm) \ = \ (1-r)b\epsilon$
$p(lm \ and \ mm \mid lm) \ = \ (1-r)b(1-\epsilon)$
$p(mm \ and \ mm \mid lm) \ = \ rb$

3) Starting from an $mm$ cell:
$p(mm \mid mm) \ = \ c$
$p(mm \ and \ mm \mid mm) \ = \ b$

To compute the LAD distribution, we will need to condition on the probability of each cell type's lineage surviving long-term and contributing to the final tumor. Set the probability that an $lw$ cell contributes to the tumor to $P$, and the probability that an $lm$ cell contributes to the tumor to $Q$. We already know the probability of an $mm$ cell contributing to the tumor is $\frac{s}{b}$.

Let's compute $P$. To get the probability of long-term survival when division leads to two non-$ww$ lineages, we will consider the probability that the $mm$ lineage produces surviving lineages regardless of what happens to the other lineage and add it to the probability that only the non-$mm$ (either $lw$ or $lm$) produces surviving lineages:

$$P \ = \ p(lw|lw)P + p(lm|lw)Q + p(mm|lw)\frac{s}{b} + \ p(lw, mm \mid lw)(P\frac{d}{b} + \frac{s}{b})$$
$$+p(lm, mm|lw)(Q\frac{d}{b} + \frac{s}{b})$$

Let's compute it in pieces

$$p(lw|lw)P + \ \ p(lw, mm \mid lw)P\frac{d}{b} = 0.5\epsilon(uc + 1 - r)P \ + \ ub\epsilon P\frac{d}{b}$$

(Note that $c = 1 - 2d - s$)

$$= 0.5\epsilon u(1 - 2d - s)P + 0.5\epsilon P - 0.5\epsilon rP + u\epsilon dP$$
$$= \epsilon P(0.5u - ud - 0.5us + 0.5 - 0.5r + ud)$$
$$= 0.5\epsilon P(u(1 - s) + (1 - r))$$

$$p(lm|lw)Q + p(lm, mm|lw)Q\frac{d}{b} \ = 0.5(1 - \epsilon)(uc + 1 - r)Q \ + \ \ ub(1 - \epsilon)Q\frac{d}{b}$$
$$= 0.5(1 - \epsilon)Q(u(1 - s) + (1 - r))$$

$$p(mm|lw)\frac{s}{b} + p(lw, mm \mid lw)\frac{s}{b} + \ p(lm, mm|lw)\frac{s}{b} = \frac{s}{b}(0.5uc + ub\epsilon + ub(1 - \epsilon))$$
$$= \frac{s}{b}u(0.5c + b)$$
$$= 0.5\frac{s}{b}u(1 + s) \quad \text{(since } c = 1 - 2b + s\text{)}$$

Putting it all together we have:

$$P = 0.5\epsilon P(u(1-s) + (1-r)) + 0.5(1-\epsilon)Q(u(1-s) + (1-r)) + 0.5\frac{s}{b}u(1+s)$$
$$= 0.5(1-r)(\epsilon P + (1-\epsilon)Q) + 0.5u((1+s)\tfrac{s}{b} + (1-s)(\epsilon P + (1-\epsilon)Q))$$

Let's compute $Q$ similarly:

$$Q = p(lw|lm)P + p(lm|lm)Q + p(mm|lm)\frac{s}{b}$$
$$+p(lw,mm|lm)(P\tfrac{d}{b} + \tfrac{s}{b}) + p(lm,mm|lm)(Q\tfrac{d}{b} + \tfrac{s}{b}) + p(mm,mm|lm)(\tfrac{s}{b} + \tfrac{s}{b}\tfrac{d}{b})$$

Again, computing in pieces:

$$p(lw|lm)P \; + \; p(lw,mm|lm)P\frac{d}{b} = 0.5c(1-r)\epsilon P + (1-r)b\epsilon P\frac{d}{b}$$
$$= \epsilon P(1-r)(0.5c + d) = 0.5\epsilon P(1-r)(1-s)$$
$$p(lm|lm)Q + p(lm,mm|lm)Q\frac{d}{b} = 0.5c(1-r)(1-\epsilon)Q + (1-r)b(1-\epsilon)Q\frac{d}{b}$$
$$= 0.5(1-\epsilon)Q(1-r)(1-s)$$
$$p(mm|lm)\tfrac{s}{b} + p(lw,mm|lm)\tfrac{s}{b} + p(lm,mm|lm)\tfrac{s}{b} + p(mm,mm|mm)(\tfrac{s}{b} + \tfrac{s}{b}\tfrac{d}{b})$$
$$= \tfrac{s}{b}(rc + (1-r)(0.5c) + (1-r)b\epsilon + (1-r)b(1-\epsilon) + rb(1 + \tfrac{d}{b})$$
$$= \tfrac{s}{b}(\,(0.5c + b)(1-r) + r(c + b + d))$$
$$= 0.5\tfrac{s}{b}(1+s)(1-r) + \tfrac{s}{b}r \;\; \text{(since } c + b + d = 1\text{)}$$

Putting it all together we have:

$$Q = 0.5\epsilon P(1-r)(1-s) + 0.5(1-\epsilon)Q(1-r)(1-s) + 0.5\frac{s}{b}(1+s)(1-r) + \frac{s}{b}r$$
$$= 0.5(1-r)((1+s)\tfrac{s}{b} + (1-s)(\epsilon P + (1-\epsilon)Q)) + r\tfrac{s}{b}$$

For convenience, we will divide everything by $\frac{s}{b}$ to get

$$P' = 0.5(1-r)(\epsilon P' + (1-\epsilon)Q') + 0.5u((1+s) + (1-s)(\epsilon P' + (1-\epsilon)Q'))$$
$$Q' = 0.5(1-r)((1+s) + (1-s)(\epsilon P' + (1-\epsilon)Q')) + r$$

For a tumor to have $LAD = k$, only one of the cells emerging at the $k^{th}$ division can divide symmetrically giving rise to two lineages that contribute to the tumor. Each of the other co-existing cells in that generation must differentiate or produce lineages that go to extinction. All these events are conditional on the original $lw$ cell contributing to the final tumor, so we need to divide all probabilities by $P$. Note that we assume the probability that a lineage survives long term but does not grow (does not contribute to the final tumor) to be negligible.

Let's the start by compiling the probabilities that starting with one cell of each type we end with a single long-term surviving lineage of each type after 1 generation:

$$p(only\ lw\ survives|lw) = p(lw|lw)P + p(lw, mm \mid lw)P\frac{d}{b} = 0.5\epsilon(u(1-s) + (1-r))P$$

This probability includes the trajectories involving 1) division without repair and with error-free replication leading to a single $lw$ cell, and 2) mutagenic repair leading to an $lm$ state that then divides either asymmetrically with error-free replication into an $lw$ cell or symmetrically with error-free translesion repair into a pair of $lw, mm$ cells with the $mm$ cell going to extinction. Everything is multiplied by $P$ because we want to consider the case that this resulting $lw$ cell survives long-term and contributes to the tumor. We had gone through the steps of that calculation in our calculation of $P$ above.

Analogously:

$$p(only\ lm\ survives|lw) = 0.5(1-\epsilon)(u(1-s) + (1-r))Q$$

$$p(only\ mm\ survives|lw) = u(0.5c + \epsilon b(1-P) + (1-\epsilon)b(1-Q))\frac{s}{b}$$

$$= u(0.5c + b(1 - \epsilon P - (1-\epsilon)Q))\frac{s}{b}$$

$$= u(0.5(1+s) - b(\epsilon P + (1-\epsilon)Q))\frac{s}{b}$$

$$= u(0.5(1+s) - s(\epsilon P' + (1-\epsilon)Q')\frac{s}{b}$$

Note that this is different from the computation of getting an $mm$ cell when computing $P$, because here we need to limit to the case when only the resulting $mm$ cell from symmetric division contributes to the tumor, which we did not do previously.

Starting with the $lm$ cell:

$$p(only\ lw\ survives|lm) = 0.5\epsilon(1-r)(1-s)P$$

$$p(only\ lm\ survives|lm) = 0.5(1-\epsilon)(1-r)(1-s)Q$$

$$p(only\ mm\ survives|lm) = r(c + 2b\frac{d}{b})\frac{s}{b} + (1-r)(0.5c + b(1 - \epsilon P - (1-\epsilon)Q))\frac{s}{b}$$

$$= r(1-s)\frac{s}{b} + (1-r)(0.5(1+s) - s(\epsilon P' + (1-\epsilon)Q'))\frac{s}{b}$$

And finally, starting with $mm$ cell:

$$p(only\ lw\ survives|mm) = 0$$

$$p(only\ lm\ survives|mm) = 0$$

$$p(only\ mm\ survives|mm) = (c + 2d)\frac{s}{b} = (1-s)\frac{s}{b}$$

Now, let's compute the probability for each cell type to give rise to two tumor-contributing lineages). Again, we need to condition on that original cell surviving:

$$p(2\ surviving\ lineages \mid lw) = ub\frac{s}{b}(\epsilon P + (1-\epsilon)Q)\frac{1}{P}$$

$$= us(\epsilon + (1-\epsilon)\frac{Q}{P}) = us(\epsilon + (1-\epsilon)\frac{Q\prime}{P\prime})$$

We are using $P'$, $Q'$ for convenience as you will see later

Note that since we are starting with an $lw$ cell, this is equivalent to the probability of $LAD = 0$

$p(LAD = 0) \ = us(\epsilon + (1-\epsilon)\frac{Q'}{P'})$

Continuing,

$$p(2\ surviving\ lineages\ |\ lm) = (1-r)b\frac{s}{b}(\epsilon P + (1-\epsilon)Q)\frac{1}{Q} + rb(\frac{s}{b})^2\frac{1}{Q}$$
$$= \ (1-r)s(1-\epsilon-\epsilon\frac{P\prime}{Q\prime}) + r\frac{s}{Q\prime}$$

$$p(2\ surviving\ lineages\ |\ mm) \ = \ b(\frac{s}{b})^2\frac{1}{s/b} = s$$

Now we can collect everything to compute the $LAD$ probability distribution

Recall, $p(LAD = 0) \ = us(\epsilon + (1-\epsilon)\frac{Q\prime}{P\prime})$

$$p(LAD = 1) = \frac{1}{P}(p(2\ surviving\ lineages|lw)p(only\ lw\ survives|lw)$$
$$+p(2\ surviving\ lineages|lm)p(only\ lm\ survives|lw)$$
$$+p(2\ surviving\ lineages|mm)p(only\ mm\ survives|lw))$$

Note that we already have all these components computed above.

Thinking about the next generation, we can think of the $LAD = 2$ scenario as having $LAD = 1$ with the surviving cell in the previous generation being our starting cell for the process. Notice that in this case that starting cell can be of any type, and we only need to additionally consider the probability of getting that cell type from the original $lw$ cell. So,

$$p(LAD = 2) = \frac{1}{P}(p(LAD = 1\ from\ lw)p(only\ lw\ survives|lw)$$
$$+p(LAD = 1\ from\ lm)p(only\ lm\ survives|lw)$$
$$+p(LAD = 1\ from\ \ mm)p(only\ mm\ survives|lw))$$

Let's compute the probabilities of $LAD = 1$ under different cell types. $p(LAD = 1\ as\ lw)$ is the same as $p(LAD = 1)$ shown above, and

$$p(LAD = 1\ from\ \ lm) = \frac{1}{Q}(p(2\ surviving\ lineages|lw)p(only\ lw\ survives|lm)$$
$$+p(2\ surviving\ lineages|lm)p(only\ lm\ survives|lm)$$
$$+p(2\ surviving\ lineages|mm)p(only\ mm\ survives|lm))$$

$$p(LAD = 1\ from\ \ mm) = \frac{b}{s}p(2\ surviving\ lineages|mm)p(only\ mm\ survives|mm)$$
$$= s(1-s)$$

And we can write,

$$p(LAD = 2) = \frac{1}{p}(p(LAD = 1\ from\ lw)p(only\ lw\ survives\ |lw)$$
$$+p(LAD = 1\ from\ \ lm)p(only\ lm\ survives|lw)$$

$$+p(LAD = 1\ from\ mm)p(only\ mm\ survives|lw))$$

And following the same logic we can get the recursion:

$$p(LAD = k\ from\ lm) = \frac{1}{Q}(p(LAD = k-1\ from\ lm)p(only\ lw\ survives|lm)$$
$$+p(LAD = k-1\ from\ lm)p(only\ lm\ survives|lm)$$
$$+p(LAD = k-1\ from\ mm)p(only\ mm\ survives|lm))$$
$$p(LAD = k\ from\ mm) = p(LAD = k-1\ from\ mm)p(only\ mm\ survives|mm)$$
$$= s(1-s)^k$$

And,

$$p(LAD = k\ from\ lw) = \frac{1}{p}(p(LAD = k-1\ from\ lw)p(only\ lw\ survives|lw)$$
$$+p(LAD = k-1\ from\ lm)p(only\ lm\ survives|lw)$$
$$+p(LAD = k-1\ from\ mm)p(only\ mm\ survives|lw))$$

This gives the analytical solution for $p(LAD = k)$

These recursions provide the full solution for the LAD distribution. Notice that none of the final equations depend on b or c as any lingering $P$ or $Q$ are divided by $P$, $Q$, or $s/b$ terms removing the dependence on $b$. This can be easily seen if you consider that $\frac{P}{Q} = \frac{P'}{Q'}$, $P/(s/b) = P'$, and $Q/(s/b) = Q'$, and neither of $P'$ or $Q'$ depend on b.

#### Matching observed LAD distribution with simulations

We consider LAD distributions derived from simulations or analytics to match a particular driver's observed distribution (as described below) if it falls within the 95% confidence interval bounds of the observations, assuming Poisson error. We only use the C3H tumors data in these analyses.

#### Inference of tumor initiation parameters from mouse sequencing data

In diploid cells, the pulse of mutagen creates lesions on both homologous chromosomes and on both DNA strands of each chromosome. Replication of one chromosome after the mutagen pulse will yield a pair of synthesized chromosomes with lesion-induced mutations on opposite strands: one chromosome with all the mutations on the Watson strand and another on the Crick strand. In case of DEN which predominantly damages thymines it means that one chromosome will have Ts on the Watson strand mutated while in another Ts on Crick strand will be mutated. Since these chromosomes segregate to different daughter cells after division of the mutagen-exposed cell, we will see Watson or Crick bias for mutations in individual chromosomes of each daughter cell. For each locus in diploid daughter cells both homologous chromosomes will have the same strand bias half of the time and the opposite strand bias half the time. In this case we would expect three possible states of asymmetry: both homologous chromosomes have thymines damaged on the Watson strand (~¼ of the genome), both homologous chromosomes have thymines damaged on Crick strand (~¼ of the genome), and homologous chromosomes have thymines damages on different strands (~½ of the genome) leading to the absence of asymmetry. Mutations are called with reference to the Watson strand meaning that these three states will look like prevalence of T>N mutations compared to A>N mutations (state 1), opposite prevalence of A>N mutations compared to T>N mutations (state 5), and similar numbers of complementary mutations (state 3), respectively (Supplementary Figure 5). This pattern could be inferred from the distribution of T>N/A>N mutations using a Hidden Markov Model.

##### Identification of mixtures of the tumors

DEN exposure leads to many independent tumors within the same mouse. Tumors are inspected, selected, and extracted manually, which could potentially lead to extraction of a mixture of two closely located tumors in one sample. If sub-tumors in this mixture have different sizes, this will lead to additional states of asymmetry in loci where sub-tumors have opposite W-C asymmetry that is not fully compensated due to size difference (states 2 and 4, Supplementary Figure 5).

To infer asymmetry states and identify mixtures of tumors we apply a Hidden Markov Model (HMM) to the mutations in each sample individually based on the Watson-Crick orientation of each mutation, allowing 5 possible hidden states (HMMas). Only T>N and A>N mutations were used as an input for this analysis. Tumors that have >5% of sites in all 5 states are classified as “mixed tumors”. In other tumors we observe one (absence of asymmetry in the whole genome), two, or three states of asymmetry. The vast majority of samples are not tumor mixtures as evident from more than 95% of mutations distributed across states 1, 3 and 5. States 1 and 3 correspond to emission probabilities ~0.9 of ~0.1 T>N vs A>N or vice versa and emission probabilities for state 5 are close to 0.5/0.5. We consider 40 (28/371 in C3H and 12/84 in CAST) tumors with more than 3 states to be admixed.

We also develop a second test to identify admixed samples for cases with a very small contribution from one of the tumors. If the size of one of the sub-tumors in the mixture is small enough, the mixture can still have only 3 states of asymmetry, as States 2 and 4 may not be distinguishable from the asymmetric states (States 1 and 3). First, to boost sensitivity we separate mutations that belong to two different sub-tumors by fitting the distribution of variant allele frequencies (VAF) in each sample as a mixture of two gaussian distributions. Mutations are attributed to one of the distributions based on a posterior probability > 0.8. We assume that a smaller sub-tumor will manifest itself as an influx of variants at fixed low variant allele frequency. It is important to note that other mechanisms, like error-free replication or symmetric tumors (LAD=0) can create a similar signal (Supplementary Figure 6). To understand whether or not low frequency variants belong to separate tumors we compare local asymmetries between high and low frequency variants. For the admixed samples, we do not expect either positive or negative correlations of local asymmetries, in contrast to the error-free replication or LAD=0 scenarios. To implement this logic, we run two HMMs with 3 states for mutations with low and high frequency separately. Samples that show non-concordant asymmetry between two subsets of mutations are assigned to the “mixed tumors” category (Supplementary Figure 6, middle panel). Non-concordant asymmetry means lack of significant overlap of HMMas states between mutations assigned to different tumors. We quantify the non-concordance by the deviation from the overlap expected based on the length of HMM asymmetry states identified in each tumor.

These two tests identify a total of 97 mixed samples (63/371 in C3H and 37/84 in CAST), which we exclude from subsequent analyses. Interestingly, a third test on admixture can be used to identify symmetric tumors (LAD=0) that have different sizes of subclones originating from the pair of sister cells after first division of a DEN-exposed cell (see below; top panel Supplementary Figure 6). Because these subclones are of different sizes, this pattern cannot be generated as a result of whole genome duplication as was recently observed for the CAROLI strain[2].

#### Identification of symmetric tumors (LAD=0)

Two daughter cells produced after division of the mutagen exposed cell would have anti-correlated patterns of lesion-induced mutations (top panel Supplementary Figure 6). In the initial study[3] some cases were described where the driver mutation was fixed before the first cell division and both daughter cells of the exposed cell contributed to the tumor. If the sizes of the subclones originated from the two daughter cells are comparable, then the resulting tumor would have no WC asymmetry in any of the chromosomes. These symmetric tumors have LAD=0 in our terminology. Formally, tumors that have less than 5% of the genome in States 1 and 5 of HMMas were attributed to this group (n=13).

If the sizes of the subclones are significantly different, we would still see asymmetry in the resulting tumor, although the amplitude of the asymmetry is expected to be lower than the tumors with higher LAD values. In this case we would expect to observe two modes of VAF distributions with each mode corresponding to a subclone. In turn, the asymmetries in each chromosome should be anti-correlated between mutations from different subclones. Similarly to the method described for identification of admixed samples, tumors that have at least two times higher than expected co-occurrence of states with opposite asymmetry (States 1 and 5) between mutations assigned to different VAF distributions when fitting a mixture of gaussians are assigned to the LAD=0 group (n=7).

Altogether, we identify 20 samples with LAD=0, all in the C3H strain.

#### Inference of the LAD

Tumors originated from the exposed cell have 0 generations between introduction of a pre-driver lesion and the emergence of the most recent common ancestor of the growing clone. Classification of these tumors was described above.

To classify the rest of the tumors, we use another mutational pattern specific for tumors induced by single pulse of DEN - multi-allelic sites. Multi-allelic sites arise due to the persistence of the lesion through several rounds of error-prone replication and incorporation of different nucleotides opposite to the lesion in each round[3]. The expected number of chromosomes with DEN-induced lesions get diluted by a factor of 2 with each subsequent division due to random segregation of chromosomes. Thus, the number of divisions between the DEN pulse and the MRCA (LAD) corresponds to the number of lesion-containing chromosomes in the tumor MRCA. Only chromosomes with lesions in MRCA could produce multiallelic mutations. In the diploid cell this dilution will create three possible states of density of multi-allelic sites corresponding to two, one or zero chromosomes containing multi-allelic sites.

These states are inferred using HMMmulti with three states on the multi-allelic sites (Supplementary Figure 7). Each cancer mutation was coded as “bi-allelic” or “multiallelic". Mutations that have more than 1 alternative variant and at least 3 reads supporting each of them were coded as "multi-allelic". Emission in each of the three states corresponds to the rate of multiallelic mutations. The three states in this model should be interpreted as zero, one, or two homologous chromosomes containing lesions.

Tumors that originated immediately after the first division of the exposed cell have 1 generation between introduction of a pre-driver lesion and the emergence of the most recent common ancestor of the growing clone (LAD=1). All chromosomes in the MRCA of these tumors carry lesions in both homologous copies, and thus the sequenced tumor would not have regions without multi-allelic sites. Tumors with <5% of the genome in State 1 are classified as LAD=1 tumors (n=121). If all four cells after the first division contribute to the resulting tumor, all of the genome would be in State 3, as both chromosomes in each homologous pair contain lesions. This was indeed the case for 61 tumors (54 C3H and 7 CAST). If, instead, only two or three out of four cells contribute to the grown tumor, some chromosomes could have fewer MAVs coming from one of the chromosomes in a homologous pair as a result of losing the cell containing fixed MAVs (Supplementary Figure 8). This would lead to an additional HMMmulti state (State 2) aggregating different scenarios of segregation and lost cells. In this case the sequence would have significant proportions of both State 2 and State 3 but not State 1. We make this observation for 60 tumors implying that they originated from two or three cells post-MRCA.

After the second division we expect half of the genome to carry lesions on one chromosome from a homologous pair, a quarter to carry lesions on both homologous chromosomes, and a quarter to be lesion-free. After the third division, the expected proportions of the genome in State 1, State 2, and State 3 would be 9/16, 6/16, and 1/16, respectively, and so on. For all tumors with LAD greater than 1, the prevalence of States 1, 2 and 3 are calculated as a proportion of genomic regions attributed to the corresponding HMMmulti state. The LAD for each tumor is then determined by minimizing the euclidean distance between proportions of the genome attributed to each State and the expected values for each LAD (Supplementary Figure 9).

#### Estimation of error-free replication rate (ε)

Rate of error-free replication (ε) could affect estimation of selection coefficients from LAD distribution. Thus we develop techniques to roughly estimate ε from VAF distribution. If we focus on the tumors with LAD=1 and all 4 progenitors after the 2nd division contributing to the tumor, we would expect most of the mutations originating from error-prone lesion replication to be shared by all the tumor cells and have VAF close to purity/2. However, error-free translesion replication followed by error-prone replication would lead to the presence of a mutation only in a subclone of the final tumor. If the first replication was error-free and the second was error-prone, the expected VAF of mutations would be around half of the VAF of shared mutations (if the size of the subclones stemming from two daughter cells of MRCA is similar, Supplementary Figure 10). Depending on the error-free and error-prone replication divisions, the non-shared mutations could be present in different subsets of cells in the tumor. The distributions of VAF in these tumors have a bi-modal shape with the high mode corresponding to shared mutations and the low mode aggregating sub-clonal mutations (Supplementary Figure 6, 10). The error-free replication on the first division would likely have the biggest impact because of the high demand of unrepaired lesions. However, some other scenarios could also contribute to the low mode because of low VAF resolution due to limited sequencing coverage.

To estimate the rate of error-free replication, we decompose the VAF distribution into a mixture of two normal distributions for each individual tumor using flexmix R-package. Only tumors with LAD=1 and all 4 progenitors after the 2nd division contributing to the tumor (n=61) are used for this analysis. VAF is calculated as 1-ref_Count/Covarege. Error-free replication rate is measured as the relative weight of the distribution component with the lower mean value. Samples that fail to fit the VAF distribution as a mixture of two normal distributions are excluded (n=5). The median estimate of error-free replication rate is 0.2 (Supplementary Figure 11).

Estimates of error-free replication rate of distinct driver mutations may differ from the average estimates because of the specificity of driver substitution type and position. For driver-specific estimates we repeat the analysis, subsetting to only mutations corresponding to the expression quantile, gene strand, and substitution type of the driver. We select only samples that have at least 500 mutations after subsetting. None of the samples with *Hras* driver mutations satisfy these criteria. For driver mutations in *Egfr* and *Braf* the estimates of error-free replication (0.18 and 0.21, respectively) are very similar to the genome-wide estimate (Supplementary Figure 12).

#### Estimation of repair rate (r)

In the studied system, lesions were introduced once by the pulse of the mutagen. Multiallelic sites are footprints of the lesions evidencing that some lesions could persist for several cell cycles unrepaired. The decrease in the proportion of multiallelic sites with each cell division controlled for the chromosome segregation reflects repair of the lesions. Repair rate was estimated as a rate of exponential decrease of the proportion of multi-allelic sites with the MRCA taking into account the HMMmulti state (Supplementary Figure 13, top). Repair rate can also vary between drivers because of the difference in the expression level, gene coding strand, and substitution type. For example, we expect genes with higher expression levels to have more efficient repair (Supplementary Figure 13).

To estimate repair rate for individual driver mutations we repeat the analysis by subsetting mutations corresponding to the expression quantile, gene strand and reference nucleotide of the driver mutation. Repair rate estimates for *Braf* (r=0.25) and *Egfr* (r=0.23) are similar to the average repair rate estimates for highly expressed genes (Supplementary Figure 14). For *Hras* the proportion of multiallelic sites is very low even in tumors with LAD=1 pointing to the high repair rate likely due to the thymine damage on the transcribed strand. This makes it impossible to estimate repair rate using this method.

#### Probability of mutagenic DNA synthesis prior to replication (u)

Two descendant cells of the exposed cell will have independent sets of mutations as they inherit lesions from different DNA strands and the probability for both of them to inherit a driver mutation is very low. However, around 3% of tumors have the initial exposed cell as MRCA and a shared driver mutation. This requires fixation of the lesion before replication.

We estimate the probability of synthesis before replication in these tumors using variant allele frequency distribution. This distribution is concentrated around 0.25 (or less depending on the sample purity) as most of the mutations are present in around half of the tumor cells. However, there is pronounced enrichment of mutations with VAF close to double of this value, especially for highly expressed genes. These mutations represent those fixed before replication (Supplementary Figure 15).

We quantified the rate of mutagenic DNA synthesis prior to replication as an excess of mutations with high VAF in sites of interest compared to non-expressed genes. Sites of interest are selected for each driver mutation independently taking into account gene expression quantile, transcription strand, and the reference nucleotide (Supplementary Table 1).

For each tumor we select a VAF cut-off that separates 5% of mutations with the highest VAFs in the non-expressed genes. Then we calculate the proportion of mutations with VAF above this cut-off in sites of interest within expressed genes. The excess of this proportion above 5% was attributed to mutagenic synthesis before replication.

#### Analysis of multiallelic sites in metastatic tumors

##### Enrichment of multiallelic variants in post-chemotherapy metastatic tumors

In the original work, lesion segregation patterns were also found in exposed human pluripotent stem cells, UV exposed human cells, and tumors with aristolochic acid exposure. Many metastases arise after treatment of the primary tumor. If metastasis growth occurs quickly after the treatment, the multiallelic sites originated due to treatment-induced DNA damage should be present in the metastatic tumors, and their distribution by chromosomes can be informative about the number of generations between treatment and metastatic expansion.

We studied multiallelic sites in the cohort of metastatic tumours provided by Hartwig Medical Foundation (HMF). Multiallelic sites were identified based on somatic variant calling using the hmftools (https://github.com/hartwigmedical/hmftools) pipeline provided by HMF. All positions that have two different alternative alleles were considered as multiallelic. Only SNVs were taken into account for this analysis.

Multiallelic sites could originate not only from lesion segregation but also due to multiple error-prone rounds of replication in the same site or as a result of sequencing errors. To select samples where multi-allelic sites likely resulted from lesion persistence, we calculate the excess of the number of multiallelic mutations above what is expected from multiple independent mutations for each sample. We select samples with at least 10-fold more multiallelic mutations above expectation for the subsequent analysis of multi-allelic sites distribution. The expected value was calculated as a square of rate of bi-allelic mutations multiplied by the length of the genome.

Pre-biopsy treatment information was obtained from metadata provided by Hartwig. We considered samples that received only one of the following treatment types: “Alkylating”, “Platinum”, or “Immunotherapy”. It is possible that the patients received other treatments outside categories.

##### Spectrum of multiallelic variants

To validate if MAVs originate from multiple independent mutations of the same site or from persistent lesions introduced by chemotherapy, we assess if the observed spectrum of MAVs is more similar to the spectrum expected based on the assumption of independent mutations or to the spectrum expected based on known chemotherapy-associated signatures.

The MAV spectrum is a set of probabilities of all possible double mutations. To simplify, we evaluate the probability of a single mutational representation of MAV spectra, e.g. $p_{MAV}(TAT>TCT)=(N_{MAV}(TAT>TCT,TAT>TTT)+N_{MAV}(TAT>TCT,TAT>TGT))/N_{MAV}$, where $N_{MAV}(TAT>TCT,TAT>TGT)$ is the corresponding number of MAV mutations and $N_{MAV}$ is the total number of MAV mutations.

The spectrum expected from independent mutations is calculated as the joint probability using probabilities of bi-allelic mutations observed in the samples, e.g. $p_{MAV}(TAT>TCT,TAT>TTT)=2\times p_{bi\text{-}allelic}(TAT>TCT)\times p_{bi\text{-}allelic}(TAT>TTT)$;

The spectrum of MAVs expected from persistent lesions introduced by treatment is calculated as a joint probability using probabilities of individual mutations in that treatment-associated signature, e.g.

$$p_{MAV}(TAT>TCT,TAT>TTT)=\sum_{sigs}F_{sig\_i}\times 2\times p_{sig\_i}(TAT>TCT)\times p_{sig\_i}(TAT>TTT),$$

where $F_{sig\_i}$ is a fraction of bi-allelic mutations coming from signature *i.*

The spectrum of observed MAVs is calculated as a proportion of each mutation type (substitution plus 3' and 5' nucleotides) counting each different substitution in the same position as a separate mutation.

The mutational spectrum of MAVs in both MAV-enriched and non-enriched samples from platinum-treated patients is dominated by T>N mutations in CTT context similar to E-SBS31 (cosmic signature SBS17) from Pich et.al[4] (Supplementary Figure 16). E-SBS31 was shown to create mutational hotspots[5] and may strongly deviate from our assumption of uniform mutation rate along the genome used in the calculation of expected numbers of multiallelic sites per sample.

We subdivide samples from platinum-treated patients into two groups based on the prevalence of E-SBS31 (threshold E-SBS31 prevalence = 10%). Samples with high prevalence of E-SBS31 show very similar patterns of multi-allelic mutations both in enriched and non-enriched samples. In contrast, the spectrum of MAVs in enriched samples is significantly different from non-enriched samples in cases with E-SBS31 < 10% (Supplementary Figure 17).

In samples with high proportion of signature E-SBS31 spectra of both MAV-enriched and non-enriched samples are very similar to the spectra expected from independent mutations in the same site (cosine similarities=1 and 0.93 correspondingly, Supplementary Figure 18). They are also consistent with the origin from E-SBS31 (cosine similarities 0.91 and 0.93).

In samples with proportion of E-SBS31 < 10%, MAV spectrum in non-enriched samples is also similar to the corresponding spectrum expected from bi-allelic mutations (cosine similarity = 0.92). In contrast, the spectrum of MAVs in MAV-enriched samples is more similar to the signature of carboplatin (E-SBS25) than to the expected spectrum from independent mutations. Interestingly, there is no footprint of E-SBS21 (SBS31 in cosmic) - the main signature associated with platinum treatment in MAVs spectrum. With this we speculate that while the main damages produced by platinum agents could not be tolerated by the replication system and can not persist for multiple cell cycles, some other damages corresponding to minor platinum signatures could lead to persistent lesions and result in multi-allelic sites.

In samples from patients treated with alkylating agents, MAVs spectra of non-enriched samples are very similar to the spectra expected under independent mutations (cosine=0.98) with a high proportion of mutations attributed to the APOBEC. In enriched samples, the observed spectrum of MAVs is not consistent with the assumption of independent mutations in the same site (cosine similarity = 0.62), nor is it consistent with known signature of cyclophosphamide, the most common medicine in the 'Alkylating' group in the dataset (Supplementary Figures 19, 20).

#### Phasing of multiallelic variant in the Hartwig cohort

Multiple mutations resulting from lesion segregation appear on the same strand, whereas independent mutation events have equal chance being on the same or different DNA strands. To phase alternative alleles of the multi-allelic variants we first select those variants that have at least one germline variant within 150 nucleotides around the somatic variant based on the germline vcf files provided by Hartwig. We then subtract mini-bam files from the original bam files containing 200nt from both sides around these variants. Using pileup, we annotate how often each allele in a multi-allelic SNV co-occurs with each of the two alleles of the heterozygous germline SNP. Cases where both alleles of SNV co-occur with the same germline variant are classified as “matching”, and cases where alternative alleles are present on different haplotypes are classified as “opposite”; ambiguous cases are excluded from the analysis (Supplementary Figure 21). The significance of the association between sample enrichment in multiallelic sites and the proportion of matching SNVs is evaluated using the Cochran–Armitage test.

We note that the null expectation for the observed fraction of somatic SNVs on the same germline haplotype is below 50%. A pair of mutations composing an MAV has a total frequency up to 50% if they occur on the same chromatid and up to 100% if they occur on different chromatids. Given that purity is far below 100%, detection of low frequency variants has limited power. This power limitation enriches for MAVs that are composed by a pair of variants on different chromatids, if these mutations accumulated independently. Additionally, most detected mutations in tumors with similar coverage are clonal[6,7], but mutations on the same chromatid have to be subclonal. Indeed, phasing data shows that for tumors not enriched in MAVs, mutations occur more often on different chromatids (n=42) compared to the same chromatid (n=18) (p-value<0.001, Chisq test).

#### LAD estimation in human metastasis

Inspired by the LAD inference performed in DEN-induced mouse liver tumors, we develop an analogous approach for human metastatic cancers. We first select metastases enriched in MAV mutations (ten-fold above expectation under independence), which are indicative of lesion segregation. To ensure sufficient power to detect LAD, we restrict the analysis to samples with at least five MAVs. For these tumors, we examine the distribution of MAVs across chromosomes, as this statistic is informative about LAD. Intuitively, if a mutational pulse occurred several divisions prior to the most recent common ancestor (MRCA) of the tumor, only a small subset of chromosomes in the MRCA cell would retain lesions. Consequently, MAVs would cluster on the lesion-bearing chromosomes.

Formally, we assume that after $n–1$ divisions (where LAD = $n$), a chromatid initially carrying lesions retains them with probability $2^{-(n-1)}$. For each chromosome $i$, there are three possible lesion states:

1. **Both chromatids retain lesions**
    - Probability: $2^{-2(n-1)}$
    - Expected MAV rate: $E(M_i) = 2u_i$
2. **One chromatid retains lesions**
    - Probability: $2×2^{-(n-1)}(1 – 2^{-(n-1)})$
    - Expected MAV rate: $E(M_i) = u_i$
3. **Neither chromatid retains lesions**
    - Probability: $(1 - 2^{-(n-1)})^2$
    - Expected MAV rate: $E(M_i) = 0$

Here, $u_i$ represents the exposure of a chromatid to lesions, which we approximate using the density of biallelic mutations on chromosome $i$.

To generate the null distribution for each LAD, we estimate $E(M_i)$ for each of 22 autosomes. We assume that, at null, MAVs are distributed across chromosomes with the same proportions. In simulations, we redistribute MAVs randomly across chromosomes with probabilities $E(M_i)/\sum E(M_i)$. This procedure was repeated 1000 times per LAD to obtain an empirical null distribution for the number of chromosomes bearing MAVs or, in other words, for MAV clustering statistics. We then compare the observed number of chromosomes containing multiallelic variants with the expected distribution for each LAD.

#### MAVs on the phylogenetic trees

Recently segregating lesions were annotated on phylogenetic trees obtained for single-cell derived blood colonies[8]. The advantage of the phylogenetic trees is that it is possible to observe not only the multiallelic variants (MAVs) but also phylogeny-violating variants (PVVs). We use these data to infer rapid expansions on the trees.

#### MRCA inference for chemotherapy treated patient

Two individuals in the analyzed cohort were chemotherapy treated patients. Patient PX002_2_01 carries only 1 MAV and 12 PVVs in total that were distributed between different clades with no more than 4 cases on the same node, so we excluded their data from subsequent analysis. Patient PX001_2_01 carries 90 MAVs and 90 PVVs. We select nodes with >=5 MAVs and apply the same method as for metastatic tumors to infer the number of divisions between exposure and clonal expansion.

#### Inference of time between treatment exposure and expansion of the clone

The logic behind this analysis is similar to the analysis of LAD in metastatic tumors. Here, we also take into account potential somatic recombination. Recombination should increase the number of chromosomes containing lesions and could thus bias the LAD estimate to smaller values. We test scenarios of 1-3 recombinations per chromosome[3,9] and simulate the expected number of chromosomes containing lesions for each LAD, accounting for length of each chromosome and its mutability (approximated from relative rate of biallelic mutations).

#### WC asymmetry anti-correlation

We calculate WC asymmetry per chromosome on a tree node as the ratio of complementary mutations attributed to this node. We restrict the analysis to T>N/A>N mutations common for the mutational signature of procarbazine. Anti-correlation means that the same chromosome displays opposite directions of WC asymmetry in two subclades.

#### PVVs in phylogeny of aging blood

Only samples from the “adult HSPCs” cohort that have multiallelic variants (n=15) are analyzed .

We claim that the ability to identify PVVs on the phylogenetic tree is informative about recent clonal expansion. We test whether the clades with PVVs demonstrate features of phylogenetic trees characteristic for clonal expansion. We perform two following tests:

1. Time in the number of mutations between the lesion node and lesion repair node is smaller than the average number of mutations between nodes with the same

topology outside the clade with PVV. We interpret this observation as the corresponding time difference.

Distribution of numbers of mutations between the lesion node and lesion repair node across all PVVs on the tree was used as an observation. We randomly select 10 control node pairs for each observed node  pair, so that they have more than 60 mutations from the root (to exclude early stages of more frequent divisions), are not terminal nodes, and are separated by the same number of divisions.

2. The number of branching nodes below the lesion node is larger than the number of branching nodes below randomly chosen control nodes. We control for the number of mutations downstream of the node and for the topology allowing for the identification of PVVs.

   We fixed the number of mutations below the node of interest equal to 200 mutations and calculated the distribution of the numbers of branching nodes spanning this interval. For the control set we have selected non-terminal nodes that have more than 60 mutations from the root (to exclude top of the overall tree with more frequent divisions), at least 2 ancestor nodes (so that we could potentially be able to identify PVV) and with 200 mutations downstream.

#### Code Availability

The code is available publicly at https://github.com/mahashady/LesionSegregation.git

### Supplementary Figures

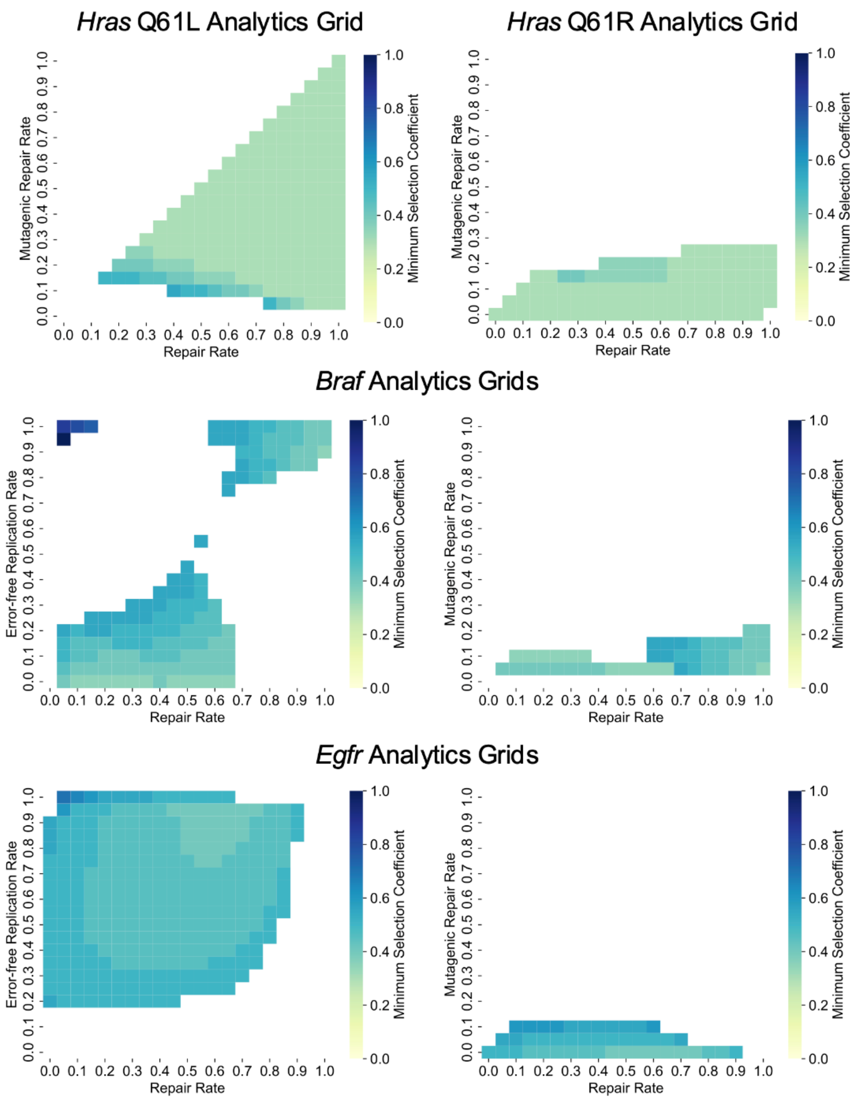


**Supplementary Figure 1**. Minimum driver selective advantage that can fit the estimated LAD distribution for the (top) *Hras* driven tumors (Q61L on the left and Q61R on the right), (middle) *Braf* driven tumors, and (bottom) *Egfr* driven tumors for all parameter combinations based on analytical calculations (Methods). For the *Hras* tumors, the y-axis of each heatmap represents the mutagenic repair rate ($u$). For the *Braf* and *Egfr* tumors, the y-axis of the left plot shows the error-free replication rate ($\epsilon$), and the y-axis of the right plot shows the mutagenic repair rate ($u$). The x-axis of all plots represents overall repair rate ($r$, mutagenic+full repair). The color at each box shows the minimum possible value of $s$ over all possible values of $u$ for a given $\epsilon, r$ pair (for $\epsilon\ vs\ r$ plots) or over all possible values of $\epsilon$ for a given $u, r$ pair (for $u\ vs\ r$ plots) that can fit the estimates for the indicated driver assuming Poisson error.

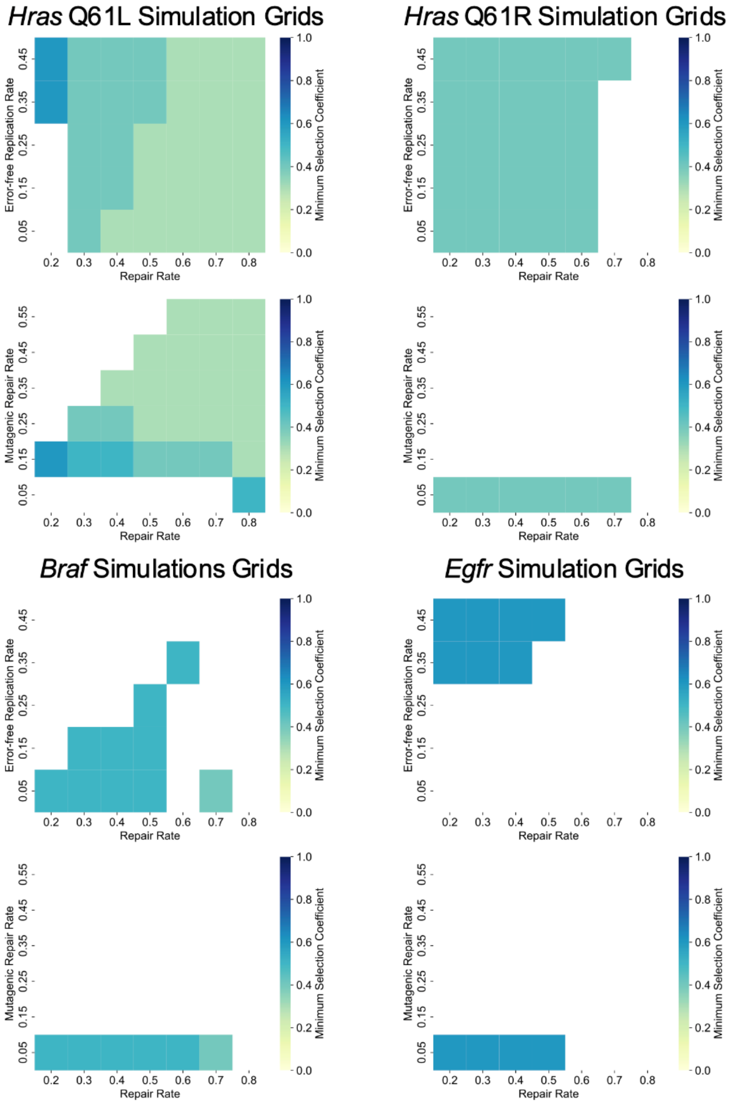


**Supplementary Figure 2**. Minimum driver selective advantage that can fit the estimated LAD distribution for the (top left) *Hras* Q61L driven tumors, (top right) *Hras* Q61R driven tumors, (middle row) *Braf* driven tumors, and (bottom row) *Egfr* driven tumors for all parameter combinations based on single driver simulations (Methods). For the *Hras* tumors, the the y-axis shows the mutagenic repair rate ($u$), and for the other drivers, the y-axis of the left plot shows the error-free replication rate ($\epsilon$) and the y-axis of the right plot shows the mutagenic repair rate ($u$). The x-axis of all plots represents overall repair rate ($r$, mutagenic+full repair). The color of each box in the heatmaps shows the minimum possible value of $s$ over all possible values of $u$ for a given $\epsilon, r$ pair (for $\epsilon\ vs\ r$ plots) or over all possible values of $\epsilon$ for a given $u, r$ pair (for $u\ vs\ r$ plots) that can fit the estimates for the indicated driver assuming Poisson error.

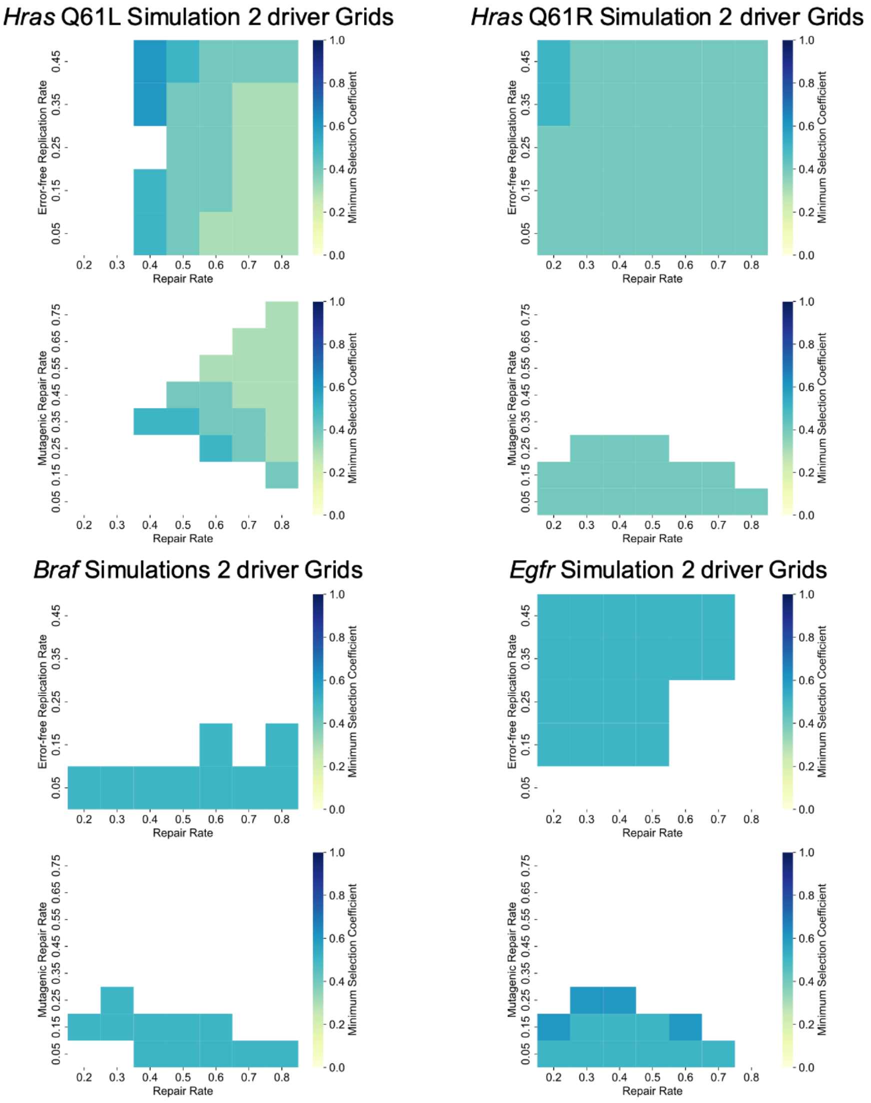


**Supplementary Figure 3**. Minimum driver selective advantage that can fit the estimated LAD distribution for the (top left) *Hras* Q61L driven tumors (top right) *Hras* Q61R driven tumors, (bottom left) *Braf* driven tumors, and (bottom right) *Egfr* driven tumors for all parameter combinations based on two driver simulations (Methods). For each driver, the y-axis of the top plot shows the error-free replication rate ($\epsilon$) and the y-axis of the bottom plot shows the mutagenic repair rate ($u$). The x-axis of all plots represents overall repair rate ($r$, mutagenic+full repair). The color at each box shows the minimum possible value of $s$ over all possible values of $u$ for a given $\epsilon, r$ pair (for $\epsilon$ $vs$ $r$ plots) or over all possible values of $\epsilon$ for a given $u, r$ pair (for $u$ $vs$ $r$ plots) that can fit the estimates for the indicated driver assuming Poisson error.

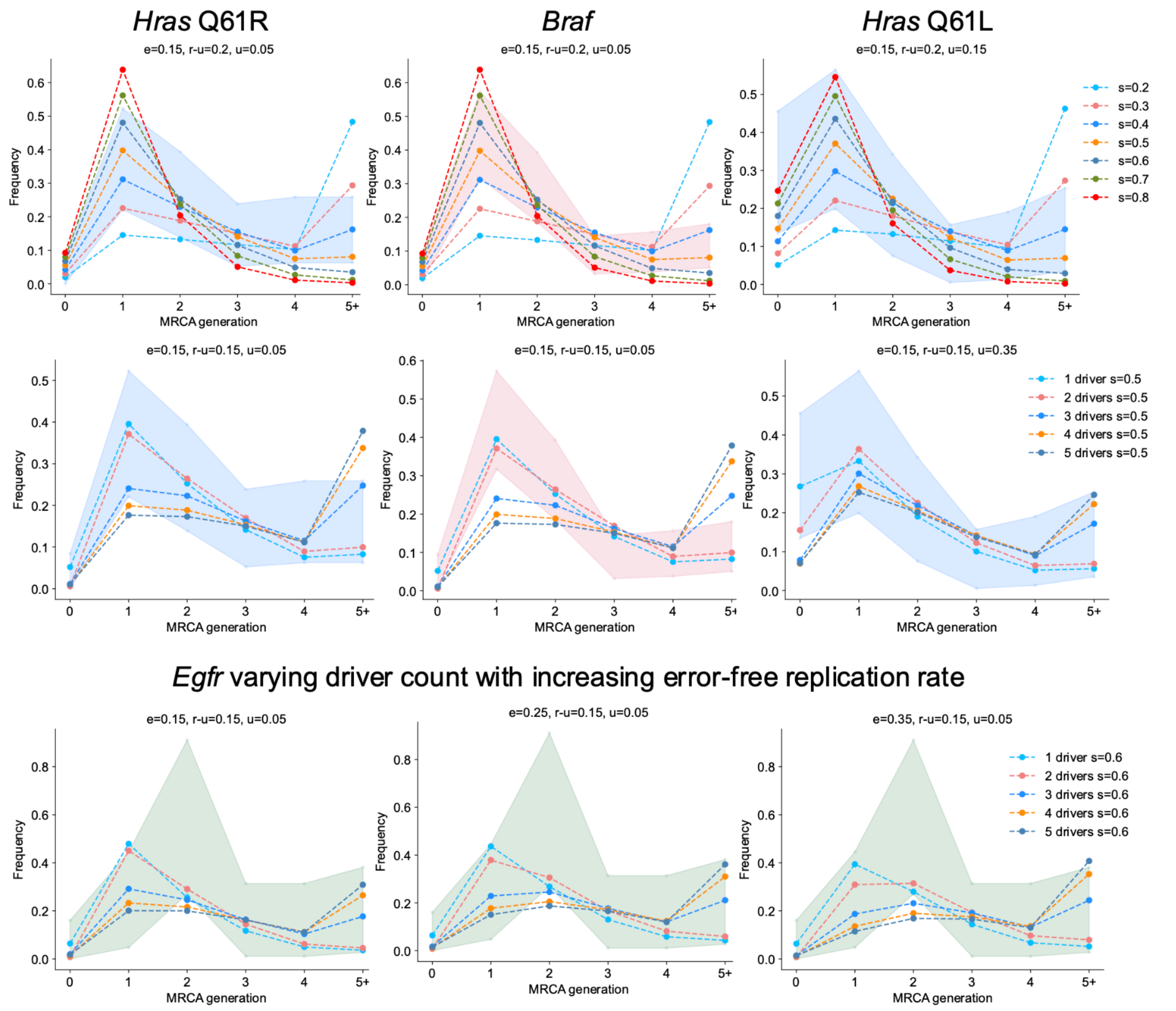


**Supplementary Figure 4.** Simulated LAD distributions under different parameter combinations (shown by lines) against estimated LAD distributions (95% Poisson confidence intervals shown by shaded background) for each driver. The top row shows the simulated distributions for selection coefficient values 0.2-0.8 for a single driver against the estimates for (left) *Hras* Q61R tumors, (middle) *Braf* V637E tumors, and (right) *Hras* Q61L tumors under non-mutagenic repair rate $(r-u)$ of 0.2, mutagenic repair $(u)$ of 0.05 for *Hras* Q61R and *Braf* and 0.15 for *Hras* Q61L, and error-free replication rate $(\epsilon)$ of 0.15. The middle row shows the simulated distributions for varying driver counts (1-5) with a final selective advantage $(s)$ of 0.5 for each driver count against the estimates for (left) *Hras* Q61R, (middle) *Braf,* (right) *Hras Q61L* driven tumors. The parameters in these plots are $r-u=$ 0.15, $u=$ 0.05 for *Hras* Q61R and *Braf* and 0.35 for *Hras* Q61L, and $\epsilon=$ 0.15. The bottom row shows the simulated LAD distributions for 1-5 drivers, all with final selective advantage $(s)$ of 0.6 under parameters $r=0.15, u=0.05$ and increasing $\epsilon$ (0.15-0.35) against the estimates for *Egfr* F254I tumors.

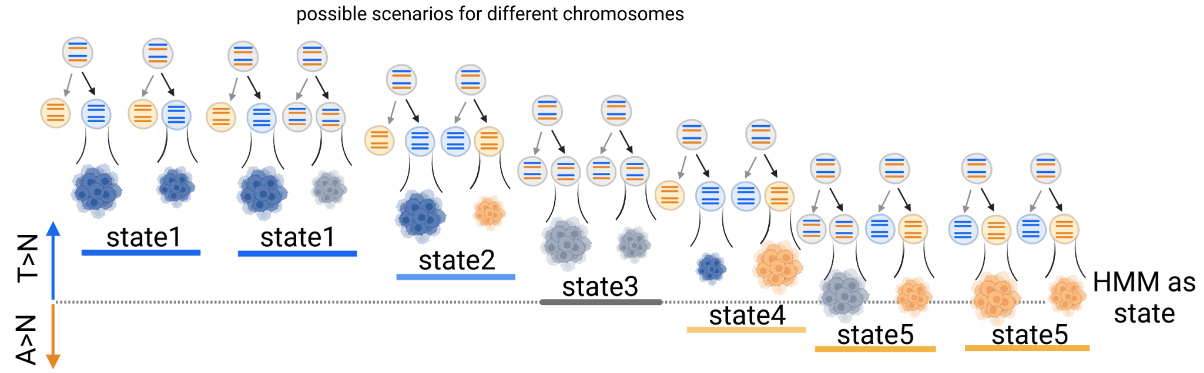


**Supplementary Figure 5.** Schematic figure representing Watson-Crick asymmetry in mixture of tumors for different possible scenarios of chromosomes segregation and tumor sizes. States 2 and 4 correspond to additional HMMas states due to mixture of tumors.

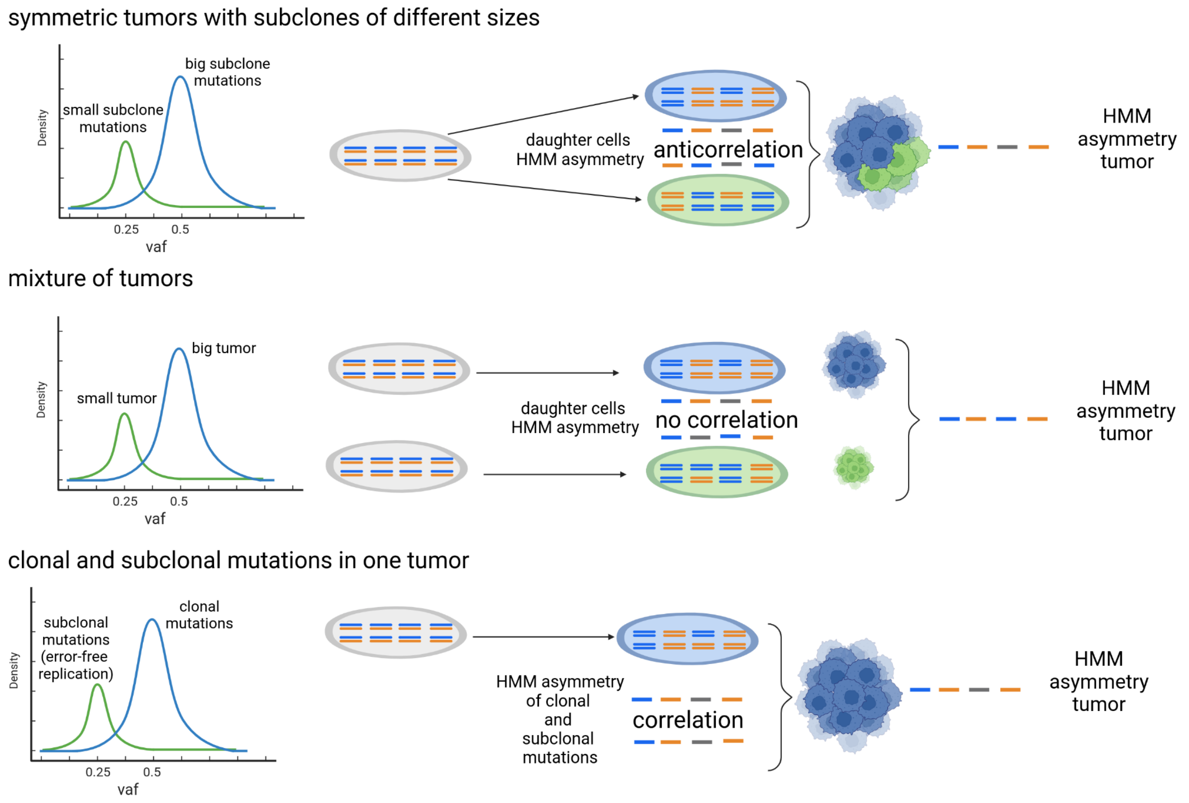


**Supplementary Figure 6.** Schematic figure representing possible explanations for bi-modal shape of variant allele frequency distribution in the sample.

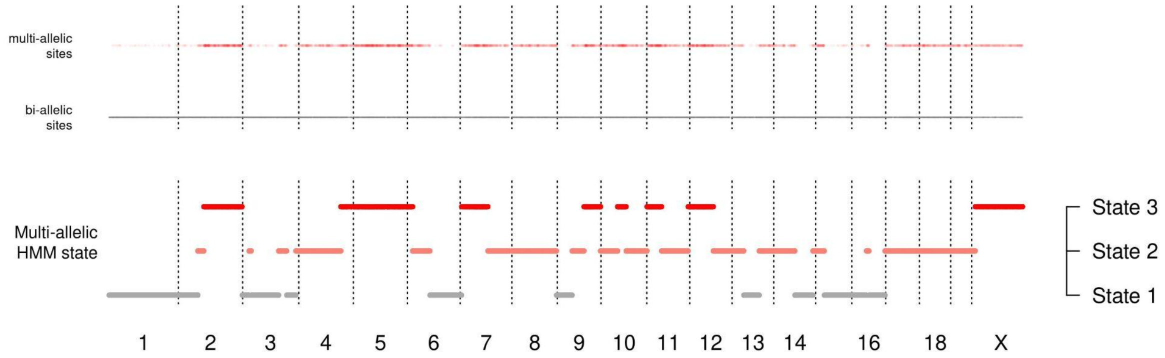


**Supplementary Figure 7.** Example of the distribution of multiallelic sites in sample 89484_N1 and states inferred by HMMmulti.

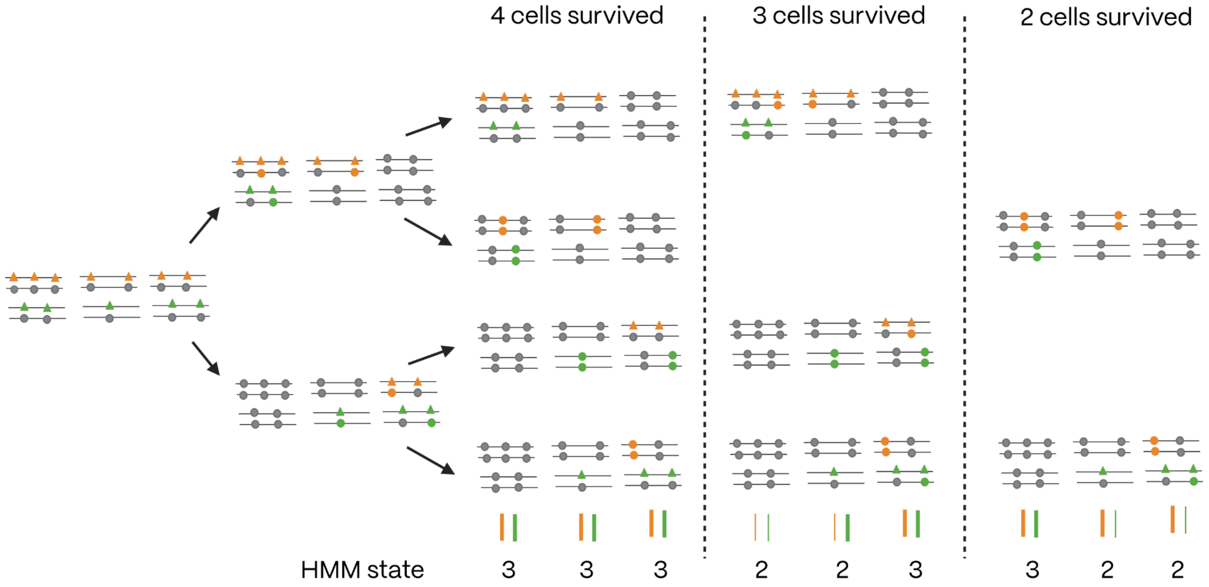


**Supplementary Figure 8.** Scheme representing states of multi-allelic HMM in case of LAD=1 and different numbers of cells survived after the two divisions. Triangles represent lesions and circles represent mutations. Different colors represent lesions and mutations on homologous chromosomes. DNA repair and error-free replication are omitted to simplify the diagram.

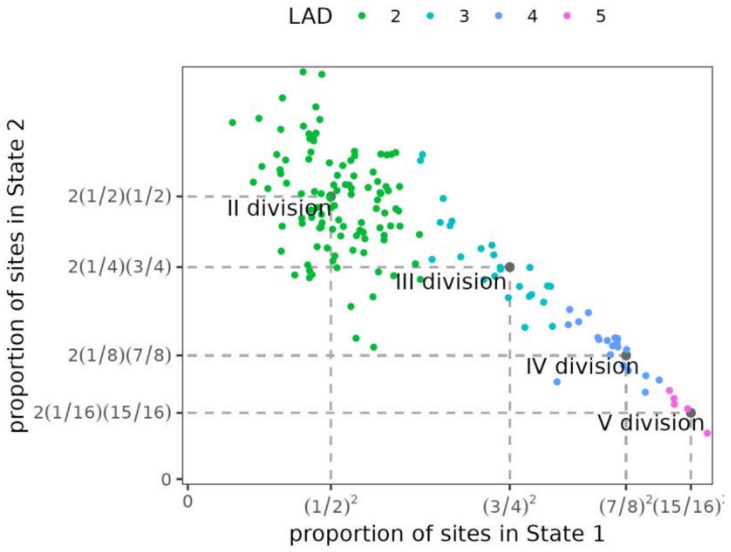


**Supplementary Figure 9.** Inference of LAD of each tumor based on the proportion of segments in different states of multiallelic HMM. Samples were attributed to different LADs minimizing euclidean distance to the expected values. Colors correspond to different LAD values.

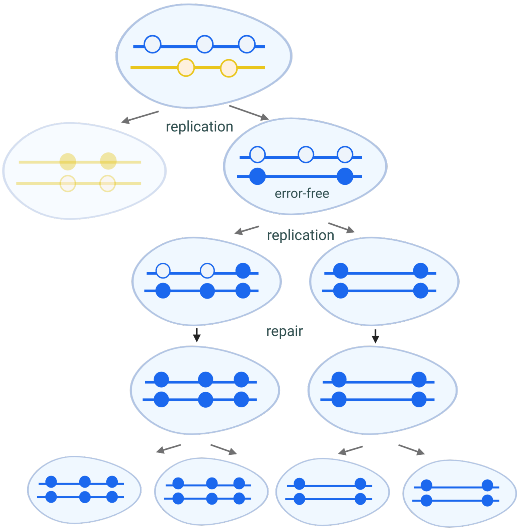


**Supplementary Figure 10.** Schematic figure showing the case where some mutations are present in only a subset of tumor cells. Cell division with error-free replication through damage results in two cells where one contains the lesion and another one has the correct nucleotide in the lesion locus. If the lesion remains unrepaired until the next replication cycle, erroneous trans-lesion replication may occur, resulting in a mutation that would be specific to descendants of this cell but not shared by all cells in the resulting tumor.

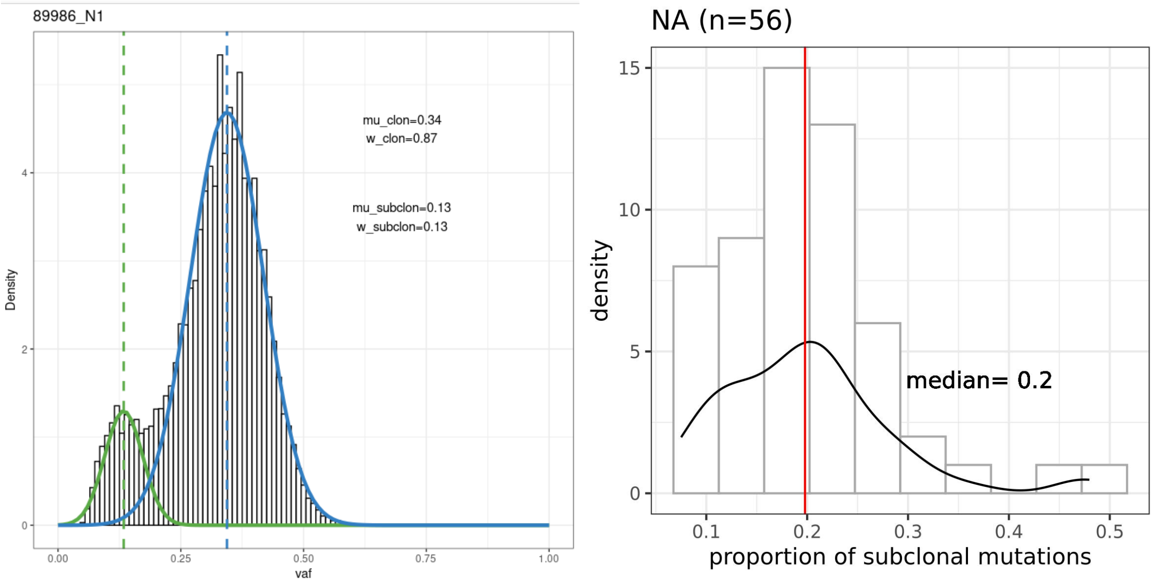


**Supplementary Figure 11.** Estimation of error-free replication rate through the proportion of mutations not shared by all the tumor cells. The left panel shows an example of a VAF distribution in one of the samples with green and blue lines showing two normal distributions resulting from the fit of the mixture of distributions. mu_clone and mu_subclone correspond to means of the resulting distributions, w_clone and w_subclone correspond to the proportion of mutations attributed to each of the resulting distributions. The right panel shows distribution of the proportion of subclonal mutations estimated as shown in the left panel in all selected samples and median of this distribution used as an aggregated value of error-free replication.

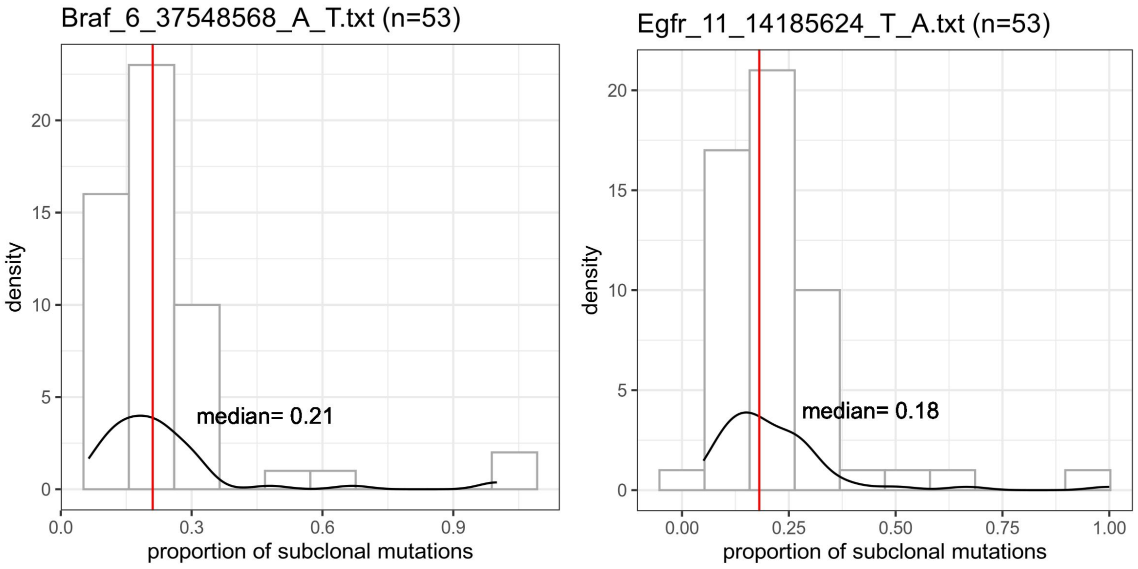


**Supplementary Figure 12.** Estimation of error-free replication rate for individual drivers.

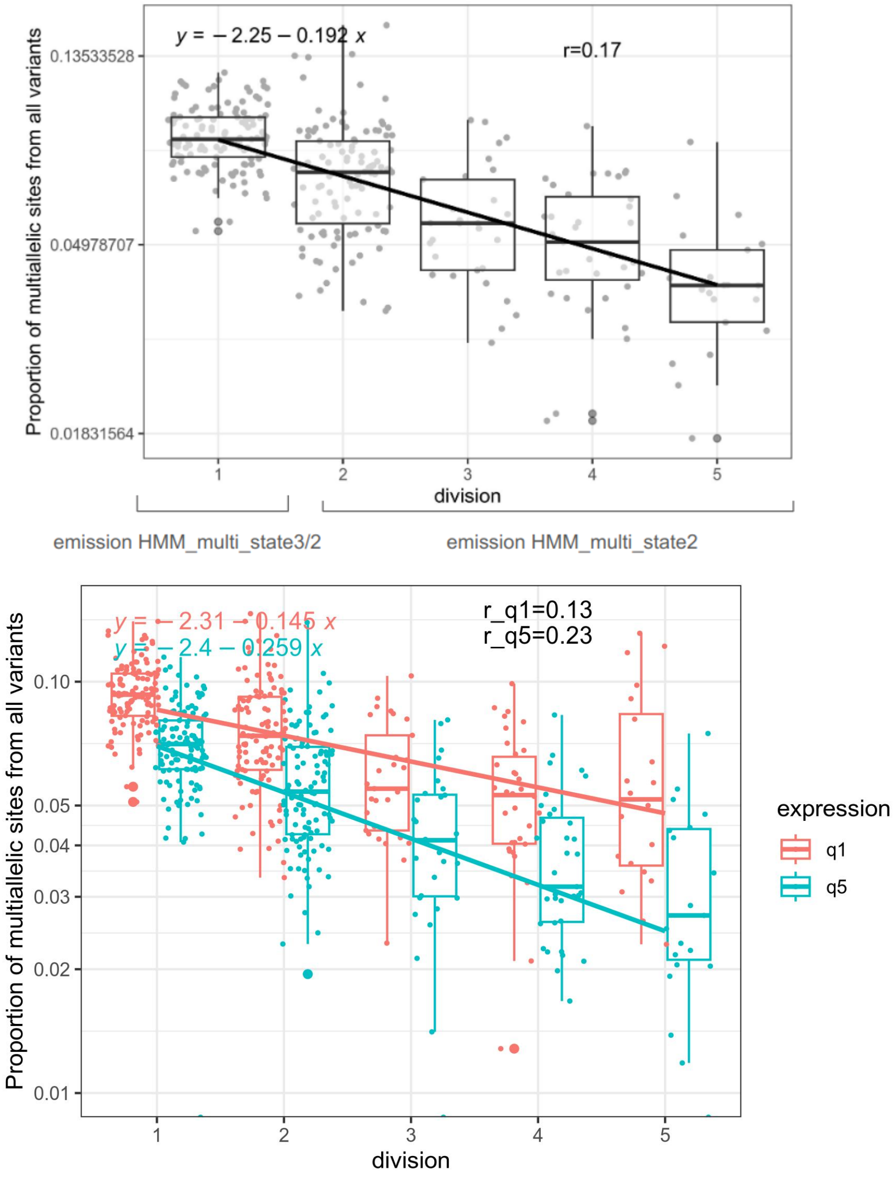


**Supplementary Note Figure 13.** Estimation of repair rate based on the decrease of the proportion of multiallelic sites with increasing MAD. Top, genome-wide estimation. Bottom, estimation based on low-expression and high-expression regions of the genome.

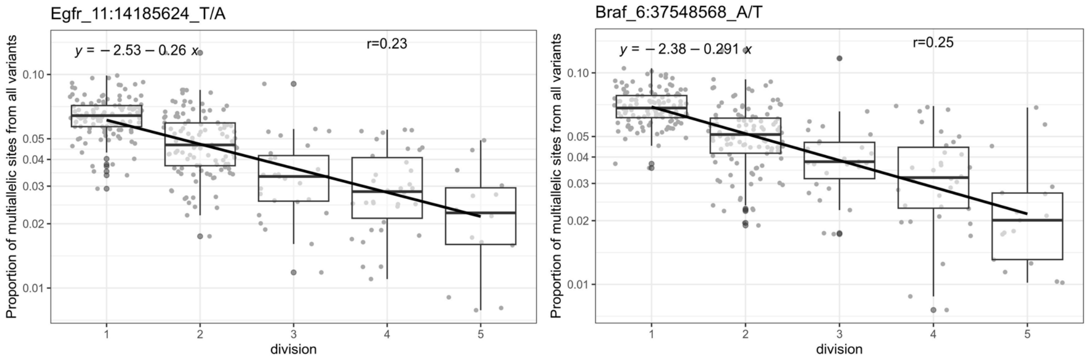


**Supplementary Figure 14.** Driver-specific estimations of repair rate.

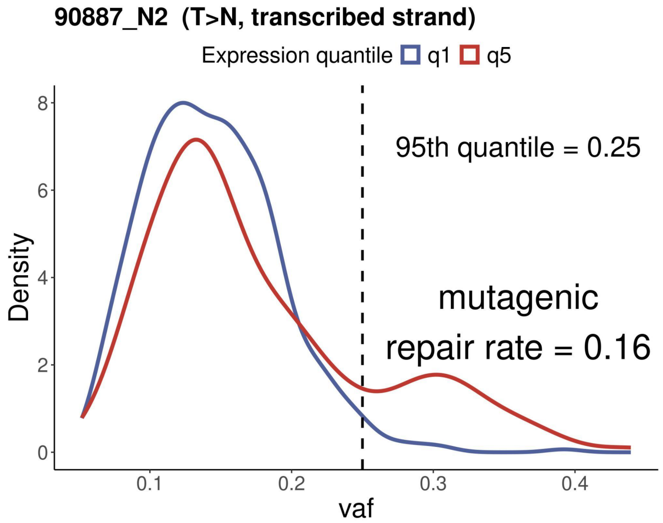


**Supplementary Figure 15.** Estimation of mutagenic DNA synthesis prior to replication based on the excess of mutations shared by all cells in the tumor originated from the exposed cell (MAD=0). Blue and red lines represent VAF distribution of mutations in low- and high-expressed genes correspondingly. 95th quantile corresponds to the line separating 5% of all mutations in the sample with the highest VAFs. Mutagenic repair rate is estimated as the excess of mutations in highly expressed genes with VAF higher than 95th quantile compared to those in low-expressed genes.

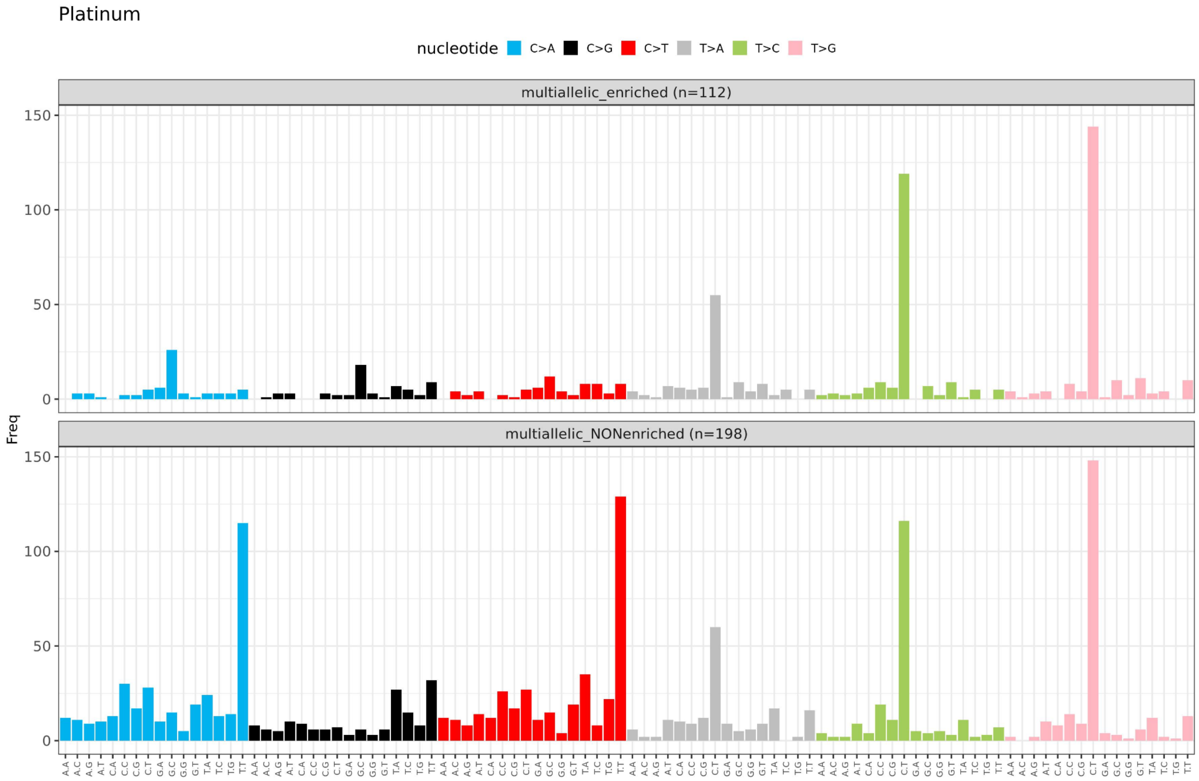


**Supplementary Figure 16.** Spectrum of multi-allelic mutations in platinum-treated patients separately for MAV-enriched and non-enriched samples. Both derived alleles contribute to the spectrum equally.

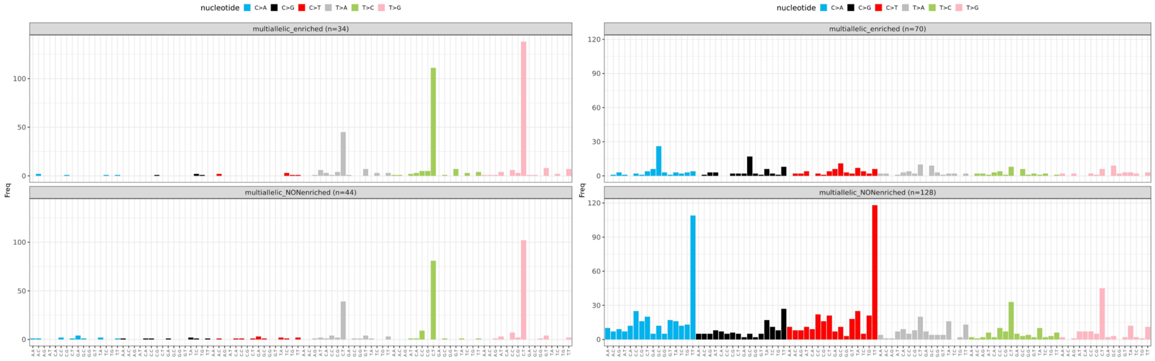


**Supplementary Figure 17.** Spectrum of MAVs in multi-allelic enriched and non-enriched platinum-treated samples with prevalence of E-SBS31 >10% (left) and E-SBS31 >10% (right).

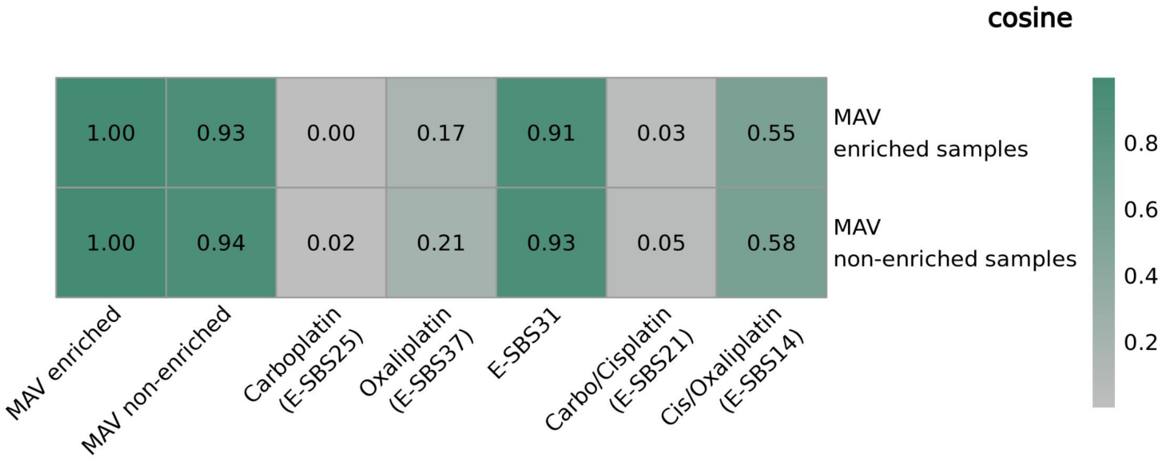


**Supplementary Figure 18.** Cosine similarities of MAVs spectra from enriched and non-enriched samples (E-SBS31 > 10%) with the spectra expected for multiple independent mutations in the same site and known spectra associated with different types of platinum treatment.

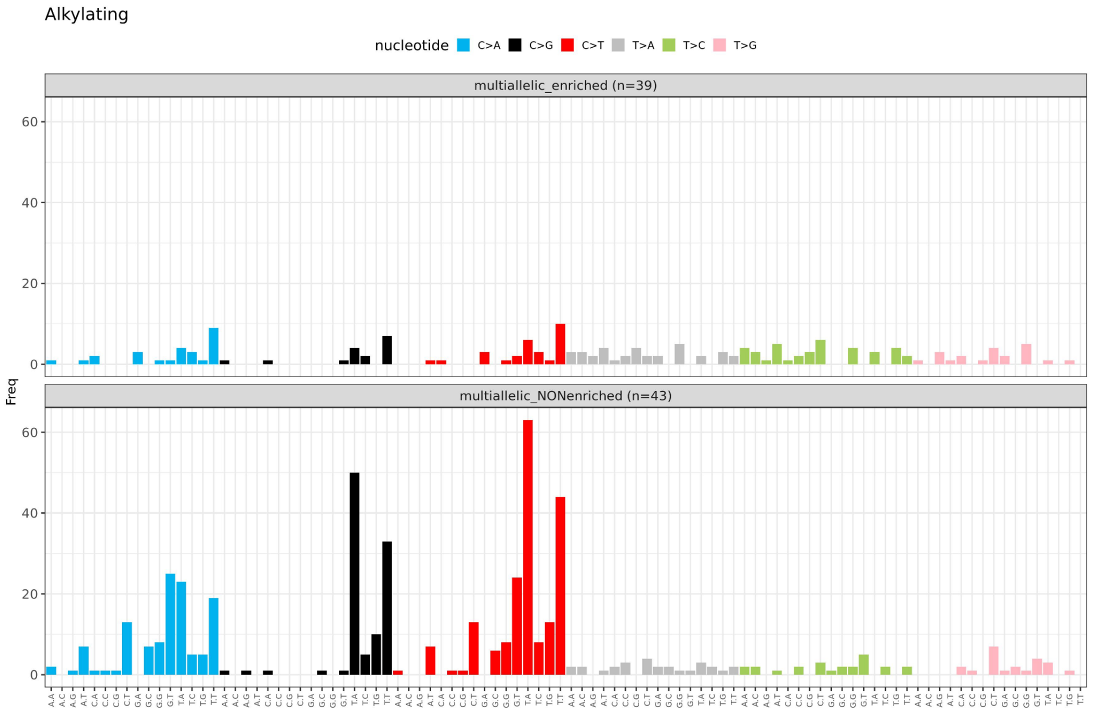


**Supplementary Figure 19.** Spectrum of MAVs in multi-allelic enriched and non-enriched alkylating-treated samples.

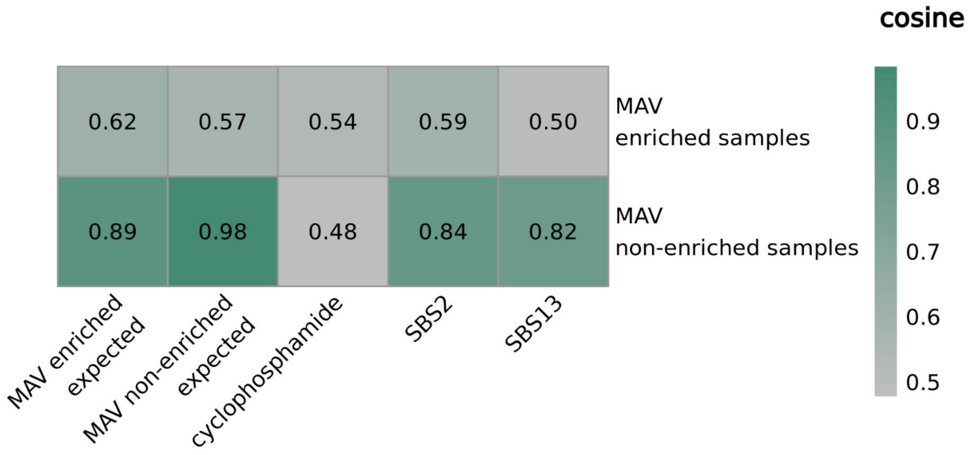


**Supplementary Figure 20.** Cosine similarity of MAVs spectra observed in MAV-enriched and non-enriched alkylating agents treated samples to the expected MAV spectra calculated based on bi-allelic mutation probabilities and known signatures of APOBEC and cyclophosphamide treatment.

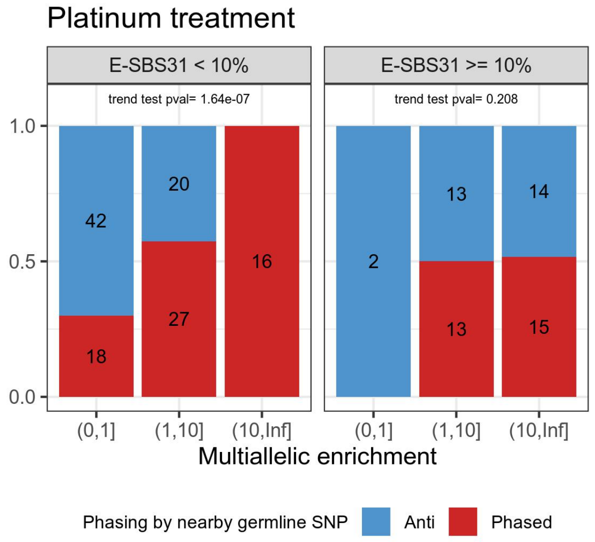


**Supplementary Figure 21 .** Phasing of MAVs with neighboring SNPs in samples with low (left) and high (right) proportion of E-SBS31. Proportions of matching and anti-pased MAVs in each category of enrichment. Numbers in the bars show absolute values of analyzed MAVs in each group. Shown p-value corresponds to Cochran–Armitage trend test.

**Supplementary Table 1.** Estimation of probability of mutagenic DNA synthesis prior to replication for individual drivers.

| Mutation | Gene | Expression quantile | Estimated mutagenic repair |
|---|---|---|---|
| 6:37548568_A/T | Braf | q4 | 0.03 |
| 7:145859242_T/C | Hras | q5 | 0.11 |
| 7:145859242_T/A | Hras | q5 | 0.11 |
| 11:14185624_T/A | Egfr | q5 | 0.04 |

**Supplementary Table 2.** Distribution of MAVs and PVVs by the nodes of phylogenetic tree contrasted based on the single cell colonies from chemotherapy treated patient.

| Node | #MAVs | #PVVs | # chroms with lesions | included in node |
|---|---|---|---|---|
| 267 | 8 | 0 | 7 | |
| 268 | 1 | 6 | 6 | |
| 304 | 6 | 0 | 5 | 267 |
| 305 | 3 | 2 | 4 | |
| 318 | 1 | 12 | 11 | |
| 342 | 6 | 0 | 6 | |
| 377 | 7 | 0 | 6 | |
| 418 | 13 | 54 | 22 | |
| 466 | 5 | 0 | 5 | |
| 492 | 11 | 0 | 8 | |

### References


1. Durrett, R. *Branching Process Models of Cancer*. (Springer International Publishing, Cham, 2015). doi:10.1007/978-3-319-16065-8.

2. Aitken, S. J. *et al.* Genetic background sets the trajectory of cancer evolution. Preprint at

https://doi.org/10.1101/2025.01.13.632787 (2025).

3. Aitken, S. J. *et al.* Pervasive lesion segregation shapes cancer genome evolution. *Nature* **583**, 265–270 (2020).

4. Pich, O. *et al.* The mutational footprints of cancer therapies. *Nat Genet* **51**, 1732–1740 (2019).

5. Arnedo-Pac, C., Muiños, F., Gonzalez-Perez, A. & Lopez-Bigas, N. Hotspot propensity across mutational processes. *Mol Syst Biol* **20**, 6–27 (2023).

6. Yan, T. *et al.* Multi-region sequencing unveils novel actionable targets and spatial heterogeneity in esophageal squamous cell carcinoma. *Nat Commun* **10**, 1670 (2019).

7. Gerlinger, M. *et al.* Intratumor Heterogeneity and Branched Evolution Revealed by Multiregion Sequencing. *New England Journal of Medicine* **366**, 883–892 (2012).

8. Spencer Chapman, M. *et al.* Prolonged persistence of mutagenic DNA lesions in somatic cells. *Nature* **638**, 729–738 (2025).

9. Ginno, P. A. *et al.* Single-mitosis dissection of acute and chronic DNA mutagenesis and repair. *Nat Genet* **56**, 913–924 (2024).
